# Preferences reveal separable valuation systems in prefrontal-limbic circuits

**DOI:** 10.1101/2023.05.10.540239

**Authors:** Frederic M. Stoll, Peter H. Rudebeck

## Abstract

Individual preferences for the flavor of different foods and fluids exert a strong influence on behavior. Most current theories posit that preferences are integrated with other state variables in orbitofrontal cortex (OFC), which is thought to derive the relative subjective value of available options to drive choice behavior. Here we report that instead of a single integrated valuation system in OFC, another separate one is centered in ventrolateral prefrontal cortex (vlPFC) in macaque monkeys. Specifically, we found that OFC and vlPFC preferentially represent outcome flavor and outcome probability, respectively, and that preferences are separately integrated into these two aspects of subjective valuation. In addition, vlPFC, but not OFC, represented the outcome probability for the two options separately, with the difference between these representations reflecting the degree of preference. Thus, there are at least two separable valuation systems that work in concert to guide choices and that both are biased by preferences.

## INTRODUCTION

Strong, trait-like preferences among foods, fluids and other primary reinforcers (or secondary reinforcers such as music, visual arts, and performance art) lead us to devote scarce resources to obtain favored options. In contrast, we may choose to forego taking a cost-free food item or ignore an artwork if we dislike it, even if it has a positive value. Research in both humans and animal models has emphasized that interaction between orbitofrontal cortex (OFC) and other parts of the limbic system, such as the amygdala, integrate preferences with an option’s costs and benefits to compute its subjective value (Gottfried et al., 2003; Schoenbaum et al., 2003; O’Doherty et al., 2006; Lopez-Persem et al., 2016). Specifically, OFC-linked brain circuits are essential for representing the features of the rewards, such as their distinct flavor or amount, that are associated with a specific choice (Schoenbaum et al., 1998; Tremblay and Schultz, 1999; Padoa-Schioppa and Assad, 2006; Stalnaker et al., 2014; Howard and Kahnt, 2017; Pastor-Bernier et al., 2019; Costa et al., 2023). Consequently, it has become widely accepted that OFC is the locus in the brain where subjective values are derived to adaptively guide behavior (Grabenhorst and Rolls, 2011; Padoa-Schioppa, 2011).

Such an OFC-centered view of how preferences and other option attributes are integrated to derive values, however, appears to be at odds with data indicating that the adjacent ventrolateral prefrontal cortex (vlPFC) plays a specific role in situations where obtaining rewards is uncertain (Murray and Rudebeck, 2018). Neural activity within vlPFC is linked to the probability of receiving rewards (Chau et al., 2015; Kaskan et al., 2017; Folloni et al., 2021; Jezzini et al., 2021) and disruption of activity within this area is associated with deficits in guiding choices based on the probability of reward (Noonan et al., 2017; Rudebeck et al., 2017; Folloni et al., 2021). Importantly, the effects seen after disruption to vlPFC are dissociable from those of OFC (Rudebeck et al., 2017), and therefore indicate that vlPFC is not simply representing integrated subjective valuations derived in OFC, but instead is representing a different aspect of subjective value. How the neural correlates of subjective value in vlPFC compare to those in OFC and how individual preferences are integrated into such representations at the level of neural activity is, however, unknown.

We reasoned that OFC and vlPFC may be the neocortical components of separable systems that work in concert with each other and with the amygdala to optimally guide choices. We hypothesized that one system, centered on OFC, preferentially represents the qualities of the outcomes (e.g., flavor) that will follow a choice, whereas another, centered on vlPFC, signals the probability that an outcome will become available (reward probability). To test this hypothesis and to establish how preferences dynamically influence these representations of value, we recorded the activity of neurons in several frontal areas and in the amygdala of macaque monkeys as they performed a probabilistic choice task for different flavors of fruit juices.

## RESULTS

### Behavioral task performance and neurophysiological recordings

We trained two rhesus monkeys to perform an instrumental choice task in which two bar-shaped stimuli appeared on each trial: one to the left and one to the right of a centrally located fixation cross (**Fig. 1a**). The height of the bar signaled the probability that juice would be delivered on that trial, and the color of the bar indicated the juice’s flavor. Following a “go” signal, monkeys were free to choose one of the two options available on that trial by fixating the corresponding response box situated next to the stimuli. A reward, the flavor and probability of which depended on the selected option, was then delivered (or not, according to a probability function). Choices from 97 and 186 sessions (from monkey M and X respectively) were analyzed using a logistic regression model that included as factors the probability associated with each outcome flavor, the previously selected outcome flavor, and an interaction term between both probabilities. Animals’ choices were strongly influenced by the probability of receiving a reward associated with each option (**Fig. 1b**). In particular, the probability associated with each of the offered outcome flavors had a significant influence on choices (at FDR-corrected p<0.05, monkey M: 95.8-97.9% of sessions; monkey X: 99.5-100% of sessions), while the interaction between probabilities had very little impact on choices (monkey M, 0%; monkey X, 9.1% of sessions).

**Fig. 1.**
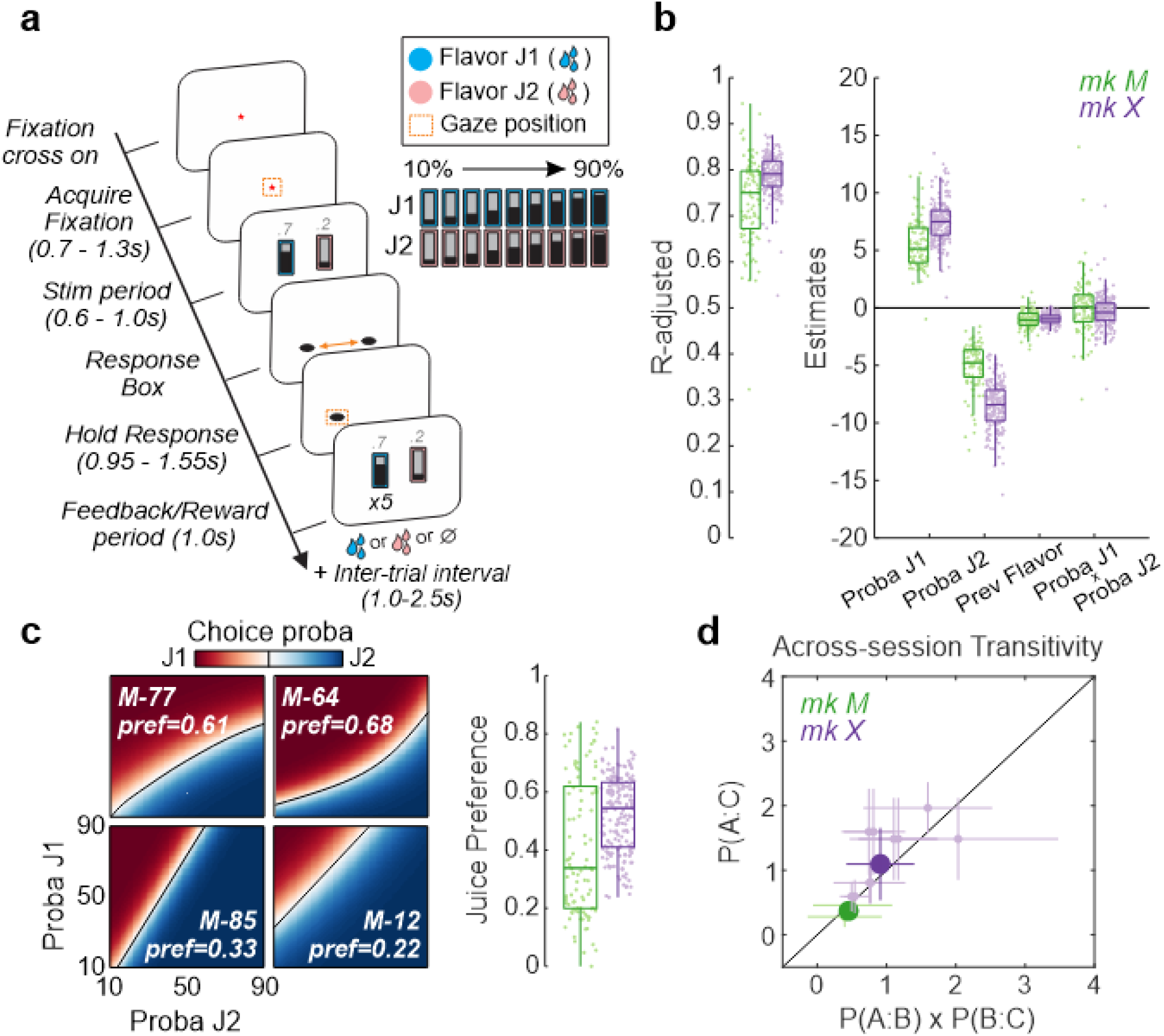
Task and behavioral performance. (**a**) In each trial, monkeys were allowed to choose one of two stimuli (Stim) presented on the screen. Stimuli comprised a background color indicating the outcome flavor and a central gauge indicating the probability of either receiving that outcome or nothing. In each session, monkeys were faced with 2 possible outcome flavors, with probabilities of reward delivery (‘proba’) ranging from 0.1 to 0.9 (or, equivalently, 10% to 90%). Monkeys made their choice with a saccade to a response box located on the side of a chosen stimulus (left or right of the central fixation cross) and then holding fixation there for the time indicated. After holding the response, we displayed the initial stimuli again, with the chosen option flashing 5 times (x5, Feedback period). Corresponding rewards were then delivered (or not) and the next trial would be initiated after an inter-trial interval. (**b**) Monkeys’ choices during each session were fitted using a logistic regression that included the probabilities associated with each outcome flavor, their interaction, and the flavor of the chosen outcome on the previous trial (*Prev Flavor*). Left and right panels show the adjusted coefficient of determination and each factor’s estimates, respectively, across sessions for monkey M (mk M, green) and monkey X (violet). Single dots represent individual sessions, with box and whisker plots indicating the median (central line), interquartile range (box), and standard deviation. (**c**) Outcome-flavor preference in four example sessions (left) and across all sessions (right) for both monkeys. A preference (‘pref’) value of 0.5 indicates no preference, whereas values close to 0 or 1 indicate a strong preference for the flavor of juice 1 (J1) or juice 2 (J2), respectively. (**d**) Measure of choice transitivity (if monkey chooses flavor A over B, and B over C, he should choose A over C) showing the odds of the choices for every triplet of flavors across sessions in both monkeys (light colors, mean ± SD), as well as the average across triplets (dark colors, mean ± SD). Values around the diagonal indicate that the monkeys’ choice across sessions followed the transitivity rule.

While the probability of receiving a reward heavily influenced subjects’ choices, monkeys’ behavior was also impacted by their individual preferences for the juices available in each session. This meant that in some testing sessions subjects were willing to choose an option with a lower probability of receiving a reward if it meant potentially receiving their preferred juice flavor. This is similar to prior observations (for example, (Padoa-Schioppa and Assad, 2006)). The relative preference for a given juice flavor – as measured by extracting the area under the modelled indifference curve between the two options (**Fig. 1c**) – varied from session to session, as different pairs of juice flavors were available in each session. Importantly, both subjects’ across-session preferences were stable and transitive; if they preferred flavor A over B, and flavor B over C, they chose flavor A over C (**Fig. 1d**, odds of choosing A > B and B > C equal A > C, two-sided sign test across monkeys, Z=-1.44, p=0.149). Thus, monkeys showed preferences for certain juices that were stable over time and transitive, confirming that they were not spurious behavioral trends but instead reflect trait-like preferences for distinct juice flavors.

### Encoding of outcome flavor and probability in frontal cortex and amygdala

Whereas lesion studies point to a role for amygdala in decisions where flavor and/or probability have to be taken into account before making a choice (Málková et al., 1997; Costa et al., 2016), there is a double dissociation of function between OFC and vlPFC (Rudebeck et al., 2017). Lesions of OFC impact decisions based on the identity of an outcome such as the flavor whereas lesions of vlPFC impact decisions based on probability. Evidence for this separation of function in frontal cortex has, however, been harder to discern at the level of neural activity.

While monkeys performed the task, we recorded the activity of 7,734 single neurons in multiple areas of frontal cortex and amygdala using a semi-chronic microdrive system (**Fig. 2a**). Recordings covered vlPFC (Walker’s area 12; n=2,714), OFC (Walker’s areas 11 and 13; n=2,556), and amygdala (AMG; n=601) (**Fig. 2b-d**). We also recorded from inferior frontal gyrus (IFG; Walker’s areas 44 and 45; n=1,008), as it was included in the prior lesion of vlPFC (Rudebeck et al., 2017), and from agranular insula (AI; lai, Iai, Iam and lapm in (Carmichael and Price, 1994); n=855), because lesions of OFC often extend into this region of posterior ventral frontal cortex. Our dataset contains neurons purely based on isolation measures (**Supplementary Fig. 1)**, with no other firing rate or selectivity restrictions. Neurons were excluded from a given analysis if the statistical model failed to converge at any time or if a subset of sessions were used based on behavioral criteria (**Supplementary Table 1**). Throughout the manuscript, only results that were seen in the activity of neurons recorded from both monkeys are highlighted.

**Fig. 2.**
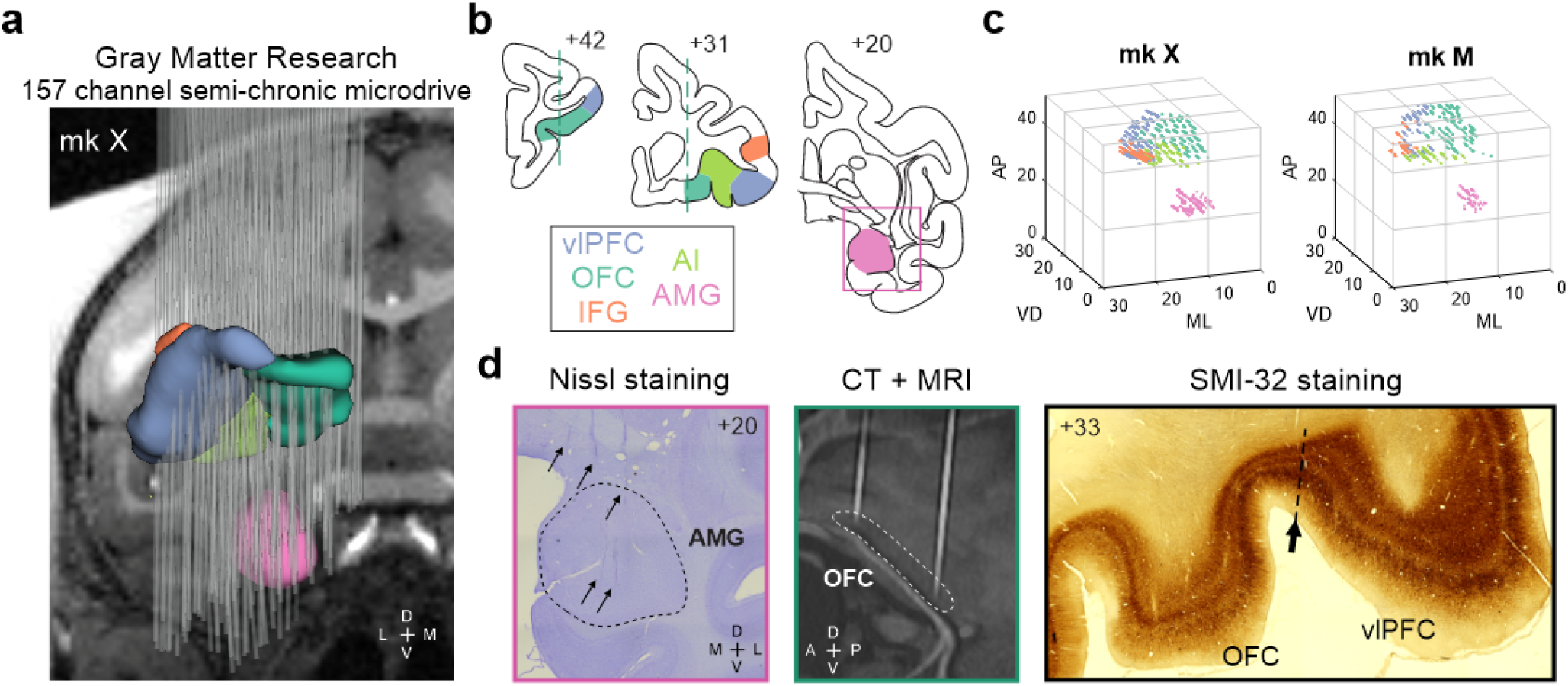
Single unit recording locations. (**a**) Model of the 157 recording electrodes (grey) and targeted areas (colors) during the MRI-based planning of the Gray Matter Research semi-chronic microdrive system used in monkey X. (**b**) Histology-based reconstruction of the brain of monkey X at different coronal sections, highlighting the targeted areas: vlPFC (blue), OFC (green), IFG (orange), AI (light green) and amygdala (pink). (**c**) 3D representation of every neuron recorded along the different electrode tracks for monkey X (left) and M (right), with colors indicating the areas assigned based on CT, MRI and histological confirmation of recording locations. (**d**) Examples of some of the methods used to confirm the precise anatomical location of recorded neurons. Left panel shows a Nissl-stained coronal section depicting the amygdala in monkey X, with arrows marking the location of 5 electrodes tracks (the pink border was reported on the corresponding coronal view on panel b). Middle panel depicts a co-registered CT and MRI in the sagittal plane, highlighting 2 electrodes tracks targeting OFC (see dashed green lines on panel b for location). Right panel shows a SMI-32 immunohistochemically stained section highlighting the boundary between OFC and vlPFC, based on the sulcus location and observation of changes in vertical processes and density of the different cortical layers as described by Carmichael and Price (1994).

We first set out to determine whether outcome flavor and probability modulated neuronal activity differentially in each area as subjects performed the task, using encoding and decoding approaches (see Methods). During the stimulus period, 30 to 40% of neurons in vlPFC, OFC, IFG and amygdala were classified as encoding the outcome flavor that would be subsequently chosen by the subject on that trial (**Fig. 3a**). By comparison, a smaller proportion of neurons in AI encoded this aspect of the task (mixed-effect logistic regression, between area comparisons in **Supplementary Table 2**). A similar pattern emerged when considering the effect size of significant neurons (mean Omega^2^ in neurons significantly representing the factor of interest during the stimulus period, **Fig. 3b**), except for OFC neurons that showed a stronger tuning compared to other areas (mixed-effect linear regression, factor area, F_(4,2348)_=6.34, p=4.5e-5, FDR-corrected post-hoc: OFC vs IFG/AI, W>2.78, p<0.018, OFC vs vlPFC/amygdala, W<1.46, p>0.20). Similarly, decoding of the outcome flavor from simultaneously recorded neurons (**Fig. 3c-d**) showed that OFC populations (1) reached higher performance across sessions compared to the other areas and (2) were more likely to significantly decode the chosen outcome flavor compared to other areas (**Supplementary Table 2**). vlPFC, amygdala (and to some extent IFG) populations reached similar decoding performance, while AI exhibited the lowest decoding performance of all recorded areas. These analyses highlight that while similar proportions of neurons in multiple areas encoded the outcome flavor that subjects chose on each trial, neurons in OFC consistently showed the highest encoding/decoding compared to all other areas recorded.

**Fig. 3.**
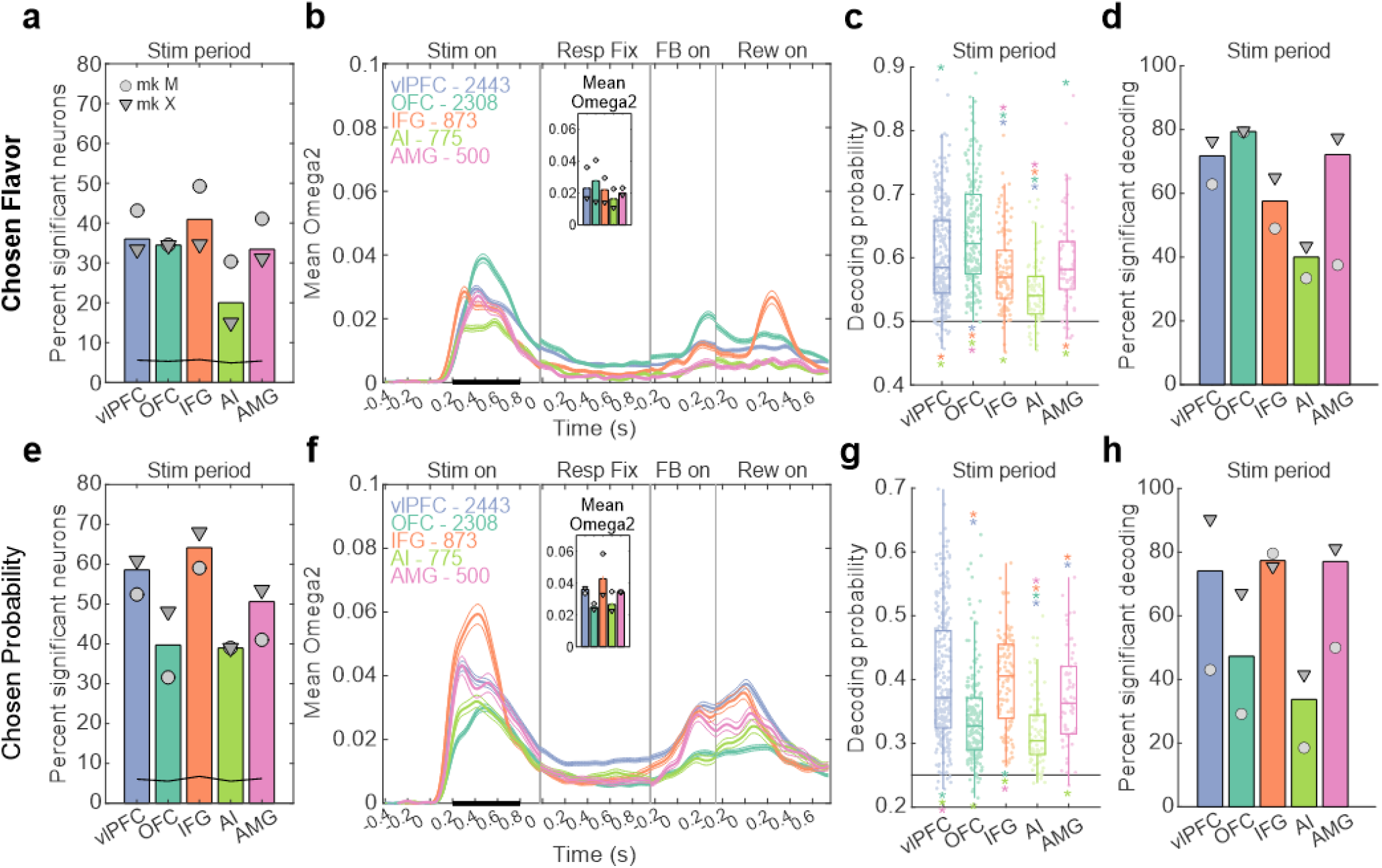
Encoding/Decoding of chosen outcome flavor (top) and chosen probability (bottom). (**a**) Percent of neurons across areas showing significant firing rate modulation with chosen outcome flavor (p<0.01 for 3 consecutive time bins) during the stimulus period (0.2 to 0.8s following stimulus onset, event ‘Stim on’). The black line shows the proportion of significantly modulated neurons during the pre-stimulus period (0.1 to 0.7s following central fixation) for each area, representing the near ∼5% chance level. (**b**) Time-resolved average effect size for neurons with significant encoding of chosen flavor during stimulus period (black bar), aligned to 4 different events (Stimulus onset, Response fixation, Feedback onset and Reward onset; each event, indicated on top, are considered time 0s and separated by vertical lines). Inset shows the average effect size during the Stim period. (**c**) Chosen flavor decoding performance using simultaneously recorded neurons in each area. Individual points represent single sessions. The statistical significances (stars) were based on a generalized mixed-effect model with area as factor and monkey/session as random intercepts (see **Supplementary Table 2**). The location of the stars (p<0.01, FDR corrected) indicate the direction of the effect (e.g. green star – OFC – above the blue boxplot – vlPFC – means that OFC decoding performance was significantly better than vlPFC). The horizontal bar represents the theoretical chance level. (**d**) Percent of sessions with a significant decoding performance, assessed using permutation testing (p<0.05). (**e-h**) As in (a-d) but for chosen probability. Color conventions as in Fig. 2. Circles and triangles represent data for monkey M and X, respectively.

Compared to the representation of outcome flavor, outcome probability elicited firing modulations in a greater proportion of neurons (**Fig. 3e-h**). Specifically, 50 to 60% of neurons in vlPFC, IFG and amygdala were classified as encoding the chosen probability during the stimulus period. A lower proportion of neurons in OFC and AI encoded this aspect of the task (**Fig. 3e** and **Supplementary Table 2**). Analysis of the mean effect sizes highlighted the prominent outcome probability encoding in neurons’ firing rate in IFG, and to a lesser extent in vlPFC and amygdala, (**Fig. 3f**; mixed-effect linear regression, factor area, F_(4,3458)_=17.7, p=2e-14, FDR-corrected post-hoc: IFG vs all areas, W>2.51, p<0.017). When we conducted a decoding analysis of outcome probability on simultaneously recorded populations of neurons, we found the most robust decoding was in vlPFC and IFG, followed by amygdala, OFC and AI (**Fig. 3g**). Notably, the proportion of sessions with statistically significant decoding performance was the highest in vlPFC, IFG and amygdala, compared to OFC or AI populations (**Fig. 3h**). We found similar pattern across areas for the representation of unchosen probability compared to chosen probability, albeit with a lower proportion of neurons significantly tuned and very little, if any, significant decoding performance (**Supplementary Fig. 2a-d** and **Supplementary Table 3**). These encoding and decoding analyses establish that outcome probability was most strongly signaled in vlPFC, IFG, and amygdala compared to OFC and AI.

To confirm that probability encoding was not simply related to response side encoding, we also determined the degree of encoding/decoding of the instrumental response (left/right) in each area. The chosen response side was most prominently represented in the activity of IFG neurons compared to other areas, with ∼70% of neurons showing firing rate modulations and high levels of decoding performance (**Supplementary Fig. 2e-h** and **Supplementary Table 3**). vlPFC and amygdala neurons only exhibited moderate encoding and decoding performance of the chosen response side, while OFC and AI neurons were relatively untuned.

Overall, these analyses complement and extend the results of prior lesion studies (Málková et al., 1997; Costa et al., 2016; Rudebeck et al., 2017) by demonstrating differential representations of flavor and probability across frontal cortex and amygdala. Notably, neuronal representations within OFC were more strongly biased toward the representation of outcome flavor compared to probability or response direction. vlPFC, IFG and amygdala populations were however more homogeneous and represented both outcome flavor and probability, although probability representations were most consistently the strongest in vlPFC and IFG. IFG population was unique in that it was most likely to represent the response direction, a feature less prominent in other areas.

### Mixed selectivity for outcome flavor and probability

Mixed selectivity at the level of single neuron activity is thought to be critical for higher level reasoning and decision-making and is prevalent within frontal cortex and amygdala (Rigotti et al., 2013; Saez et al., 2015). Although we found clear differences in the representations of the chosen flavor and probability across areas, most of the neurons that we recorded exhibited multidimensional selectivity during the stimulus period, representing both outcome flavor and probability (**Fig. 4a-b** and **Supplementary Fig. 3a**).

**Fig. 4.**
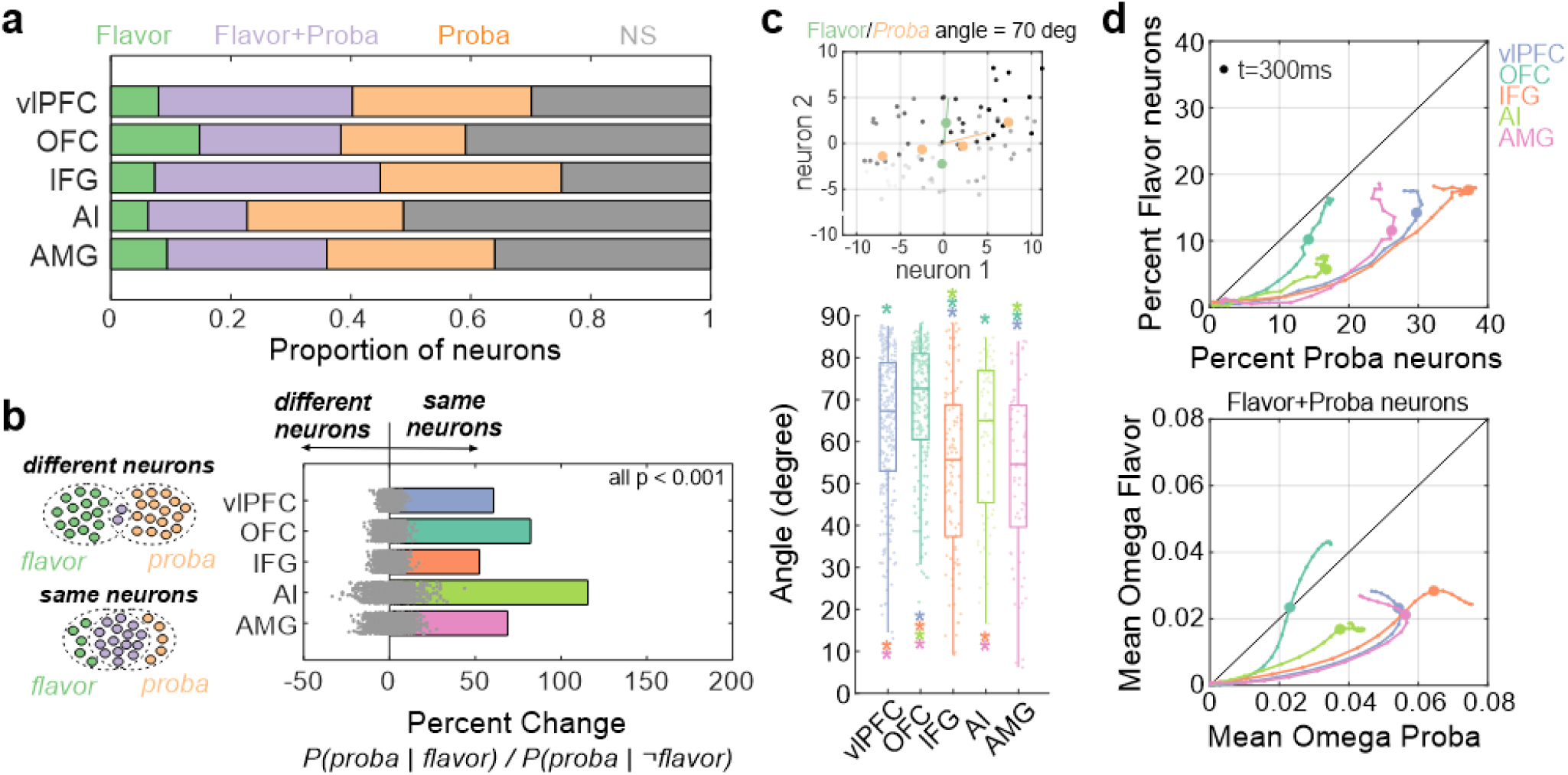
Mixed selectivity in chosen Flavor and Probability representation. (**a**) Proportion of neurons in each area showing significant encoding of either chosen flavor (green), chosen probability (orange), both (purple) or neither (grey). (**b**) Estimate of the proportion of chosen flavor neurons also representing chosen probability. Positive values indicate that the same neurons are more likely to encode both parameters, while negative values indicate that different neurons encode the two parameters. Grey dots represent the ratio extracted for each of the 1000 permutations, used to derive significance (all areas p<0.001). (**c**) Top panel shows a simulated example of the extracted angle between outcome flavor and probability subspaces, when considering only 2 neurons. Grey dots represent the normalized and centered firing rate of the two neurons across the 10 repetitions of each of the 8 conditions (2 chosen flavor x 4 chosen probability) while the colored dots represent the average across repetitions (green: chosen flavors, orange: chosen probabilities). The lines represent the 2 subspaces extracted using cross-validated PCA, from which the angle between them was extracted. Bottom panel shows the angle (between 0 and 90 degrees) between outcome flavor and probability subspaces for every simultaneously recorded neuron population (dots) and for each area, highlighting a significant difference between areas (mixed-effect linear regression, factor area, F_(4,677)_=16.3, p=8.8e-13). The location of the stars (p<0.01, FDR corrected) indicate the direction of the effect, as in Fig. 3c. (**d**) Top panel shows the evolution over time of the percent of neurons significantly encoding chosen flavor (y-axis) compared to chosen probability (x-axis) for each area. Bottom panel shows the same evolution but for the mean effect size of the significant neurons. The line and dots represent the time (larger dots at 300ms following stimulus onset), displayed from stimulus onset and until reaching the area’s maximum value in either axis.

The representation of outcome flavor and probability could nevertheless be encoded by distinct patterns of neural activity at the population level. To assess this, we computed the angle between population subspaces for outcome flavor and probability (see Methods). An angle of 0 degree means that the representations are entirely overlapping whereas an angle of 90 degrees indicates that the representations are orthogonal and largely separate (**Fig. 4c**). We found clear differences among areas, with OFC populations showing the greatest angle between outcome probability and flavor subspaces (median of 73.2 degrees; FDR-corrected post-hoc: OFC vs all areas, W>3.14, p<0.003), followed by vlPFC and AI populations (∼66.5 degrees; vlPFC/AI vs IFG/amygdala, W>2.44, p<0.018). Thus, even though the activity of single neurons in these areas was likely to represent both outcome flavor and probability, both types of information were represented within largely separate neural spaces. In IFG and amygdala by comparison, the angle between outcome flavor and probability subspaces was lower at around 55.5 degrees. This indicates that representations of outcome flavor and probability were spanning a more similar neural space in these areas, potentially indicative of these representations being more closely linked to more general processes such as salience.

Next, we looked at the variation in the timing of encoding across areas. We did this to probe how outcome flavor and probability are dynamically signaled across frontal cortex and amygdala. Here we compared the time-resolved proportion and effect size (**Fig. 4d** top and bottom, respectively, and **Supplementary Fig. 3b-c**) of neurons encoding outcome flavor and probability starting from the presentation of the two stimuli. We found an initial bias toward the representation of chosen probability in all areas, but OFC and AI populations exhibited a different trajectory compared to other areas. In these areas, the population representation quickly moved back to the identity line as the representation of probability plateaued and there was an increase in the representation of flavor. Additionally, the time at which flavor and probability information were encoded varied across areas. IFG neurons represented chosen probability earlier than all areas, while OFC and AI neurons were the latest to do so (mixed-effect linear regression, factor area, F_(4,3703)_=29.2, p=5.7e-24; FDR-corrected post-hoc: IFG vs all others, W>3.26, p<1.8e-3; OFC/AI vs all others, W>2.4, p<0.02; OFC vs AI, W=1.79, p=0.08). A similar pattern was found for the encoding of the chosen outcome flavor where IFG neurons were the earliest to do so and OFC neurons the latest (mixed-effect linear regression, factor area, F_(4,2568)_=7.61, p=4.2e-6; FDR-corrected post-hoc: IFG vs vlPFC, W>2.76, p=0.015; OFC vs vlPFC/IFG/amygdala, W>3.1, p<0.007; OFC vs AI, W=1.41, p=0.22).

Taken together, neurons in all areas exhibited multidimensional selectivity, but this selectivity was not uniform. Chosen probability encoding emerged earlier and dominated activity in vlPFC and IFG neurons compared to outcome flavor. Representations in IFG and amygdala neurons existed within a more shared neural space. Activity in OFC, in contrast, represented outcome flavor independently of probability representations, spanning different neural spaces, with this encoding emerging later than other areas.

### Session-level effects of outcome flavor preferences

Given clear evidence of session-by-session difference in preferences for the different juices (**Fig. 1c**), we investigated how the degree of preference that subjects expressed in each session impacted the ability to decode outcome flavor or probability from neural activity. One possibility is that decoding performance for outcome flavor might increase in sessions where subjects exhibit a strong preference for one of the two juices while probability decoding would not be affected (Option a: flavor specific, **Fig. 5a**). If such a pattern were observed, it would indicate that preferences bias representations of juice flavor prior to being integrated with representations of probability. Alternatively, decoding performance of juice flavor in an area could be increased in sessions where subjects exhibit a preference for one of the juices while probability decoding would be decreased (Option b: flavor interaction, **Fig. 5a**). Such a pattern would mirror the changes in behavior where subjects are less sensitive to probability in sessions with a strong preference. It would also indicate that preferences are integrated into representations of both outcome flavor and outcome probability to guide choice. Another possibility is that preferences have no impact on decoding of either outcome flavor or probability (Option c: independent, **Fig. 5a**) as they are not integrated into the decision process in that area.

**Fig. 5.**
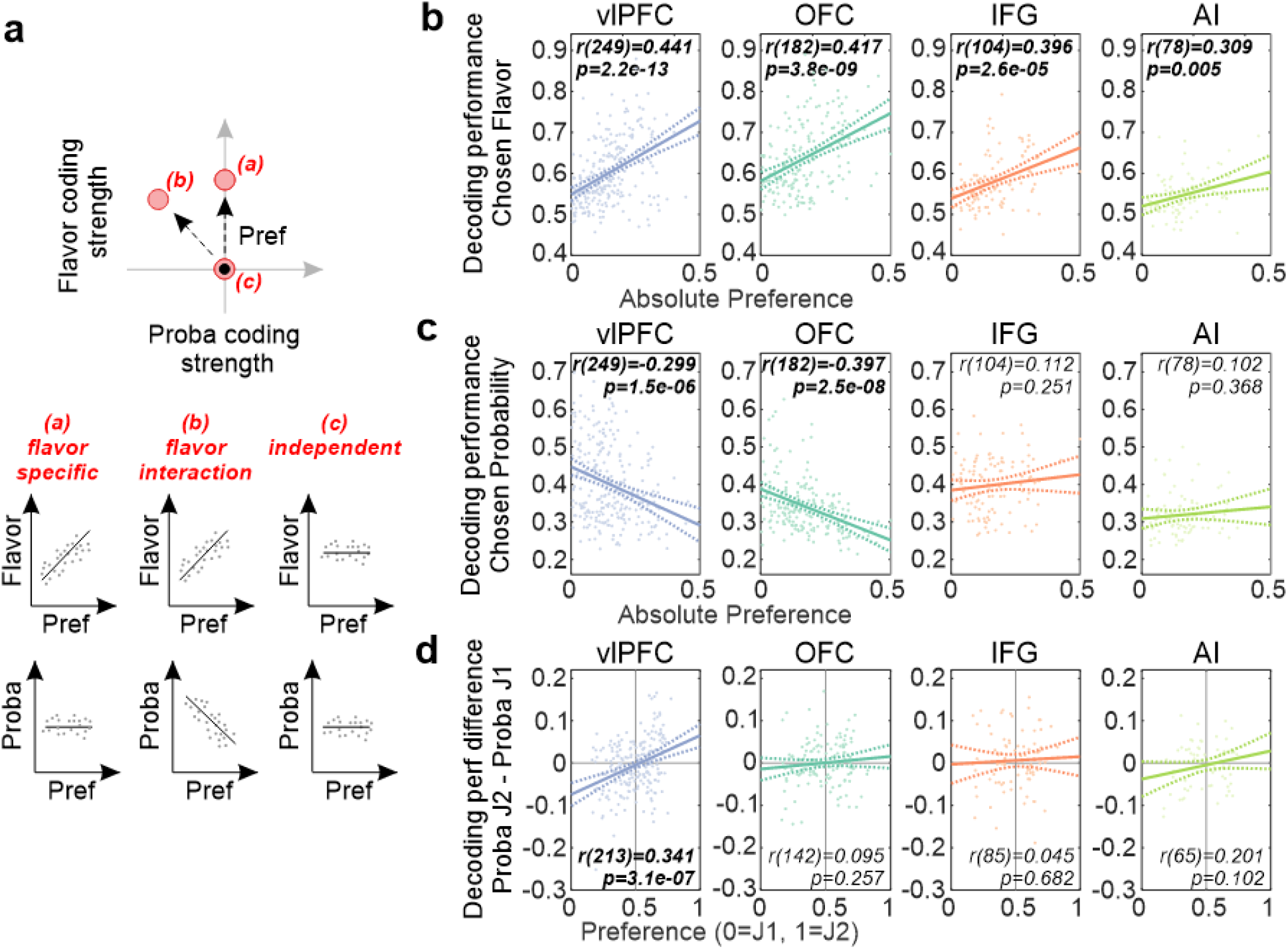
Chosen flavor and probability representations relate to outcome preference. (**a**) Schema representing the different way preference might influence the representation of chosen flavor and probability within an area. Top panel shows the 3 possible trajectories in the strength of flavor and probability coding within a neural population when subjects exhibit a strong juice preference (red dots), compared to the relative representation when they show no preference (black dot). Bottom panels show the expected correlation between decoding performance of flavor (top) and probability (bottom) with preference (as shown in panel b-c) for the 3 proposed hypotheses (flavor specific, flavor interaction or independent). (**b**) Correlation between the decoding performance for outcome flavor across neuronal populations and the absolute outcome flavor preference during the corresponding session (0 = no preference, 0.5 = maximal preference). A significant effect of preference was found (mixed-effect linear regression, factor abs(preference), F_(1,613)_=65.5, p=3.1e-15) across all areas (interaction abs(preference) x area, F_(3,613)_=1.76, p=0.15) (**c**) Same as panel b but for the decoding performance for chosen probability (mixed-effect linear regression, interaction abs(preference) x area, F_(3,613)_=9.4, p=4.4e-6) (**d**) Correlation between the difference in probability decoding performance when J1 or J2 was chosen and the outcome preference (mixed-effect linear regression, interaction preference x area, F_(3,505)_=3.69, p=0.012; note the lower degree of freedom due to further trial number restrictions).

To test among these alternatives, we assessed the relationship between the degree of juice preference across sessions (area under the modelled indifference curve) and the performance of linear classifiers described previously. Importantly, our results could not simply be explained by differences in the number of trials per category as these were matched when we constructed each one of our classifiers (see Methods). Note that amygdala recordings were not included in further analyses due to the limited number of simultaneously recorded neurons and sessions where strong preferences were exhibited.

A preference for one of the juices had a strong influence on decoding of outcome flavor and probability across sessions (**Fig. 5b-c**, **Supplementary Fig. 4-5**). First, we found that decoding of chosen flavor was higher when monkeys had stronger juice preferences (**Fig. 5b**). This was true for all areas, with the strongest modulations in vlPFC, IFG and OFC. Notably, these modulations were most consistent across monkeys in vlPFC and IFG (**Supplementary Fig. 5**). Second, probability representations in vlPFC and OFC were also correlated with juice preference, where stronger preference was associated with lower decoding performance of chosen probability (**Fig. 5c**). This effect was the most reliable across monkeys in OFC (**Supplementary Fig. 5**), indicating that preferences specifically caused a decrease in probability representations in this part of frontal cortex.

The above analyses on probability decoding were, however, performed independently of the chosen flavor. It is possible that the same probability associated with the two outcome flavors might not be represented similarly when monkeys exhibit a preference for one of the juices. If an area is integrating flavor preferences with probability separately for the two juices, then this would indicate that it is fully integrating the preferences into the subjective valuations of the different options. Consequently, we separately determined the decoding performance for chosen probability for each of the two outcome flavors. We then took the difference in performance between these two decoding measures and compared this to subjects’ preferences (**Fig. 5d**). Here, we found that only vlPFC showed a significant influence of preference on the difference in probability decoding performance, where the chosen probability of the preferred juice outcome was better represented than the chosen probability of the alternative. No other area exhibited a similar effect. Control analyses confirmed that these effects were specific; preferences did not impact decoding of the unchosen probability or the chosen response side (**Supplementary Fig. 4**).

In summary, when subjects exhibited a preference for one of the juices, this altered representations of outcome flavor and probability in frontal cortex (Options a and b in **Fig. 5**). The pattern was not, however, uniform across areas. Notably, vlPFC was unique among frontal areas as representations of probability for the two juice flavors were separable, and the difference between these representations was related to the degree of preference. This pattern of results indicates that contrary to current models of economic choice (Padoa-Schioppa, 2011), which emphasize the role of OFC, neurons in vlPFC may be a central point of integration of different aspects that are used to guide decisions, a point we take up later.

### Behavior related to within session changes in outcome flavor preference

The prior analyses establish that stable trait-like preferences for certain outcome flavors at the level of whole sessions bias choices and neural activity. While preferences were apparent at this temporal scale, it was also the case that preferences for the two juices often fluctuated within a session. This indicates that subjects were constantly updating the value of juices as they progressed through the task. One indication of within-session changes in preference was that we could observe a drift in the residuals in our initial behavioral model. A trend analysis of these residuals revealed that such a drift was present in at least 26.5% of sessions (monkey M: n=18/97; monkey X: n=57/186, MK test, FDR-corrected p<0.05). Thus, factors other than the options available on each trial influenced subjects’ choices.

To test whether this variation in the residual variance was related to a change of preference for a given outcome flavor across time, we divided each session into 50 bins of equal trial numbers using a sliding window within which we then applied our behavioral model to explain subjects’ choices (using the same model as in **Fig. 1b**). This allowed us to extract the local juice preference across the whole session (see example session in **Fig. 6a**). We found that the strength of the change in smoothed residuals correlated with the average deviation in the local juice preference across sessions (**Fig. 6b**; note that this is a negative correlation as the residuals are the inverse). Importantly, this effect was unique to local preference; other trial-by-trial measures indicative of motivation such as initiation and reaction times (log(IT) and log(RT)) were not as correlated with the residuals or local preferences (**Supplementary Fig. 6**). This was also evident at the level of individual sessions (**Fig. 6c**). Notably, a similar pattern of effects was observed when assessing performance in a selective satiety test in one animal (Monkey X, n=6 sessions, **Supplementary Fig. 7**). Taken together, this indicates that the within-session variation in preference for the different juices were goal-directed and not related to aspects of inattention or fatigue.

**Fig. 6.**
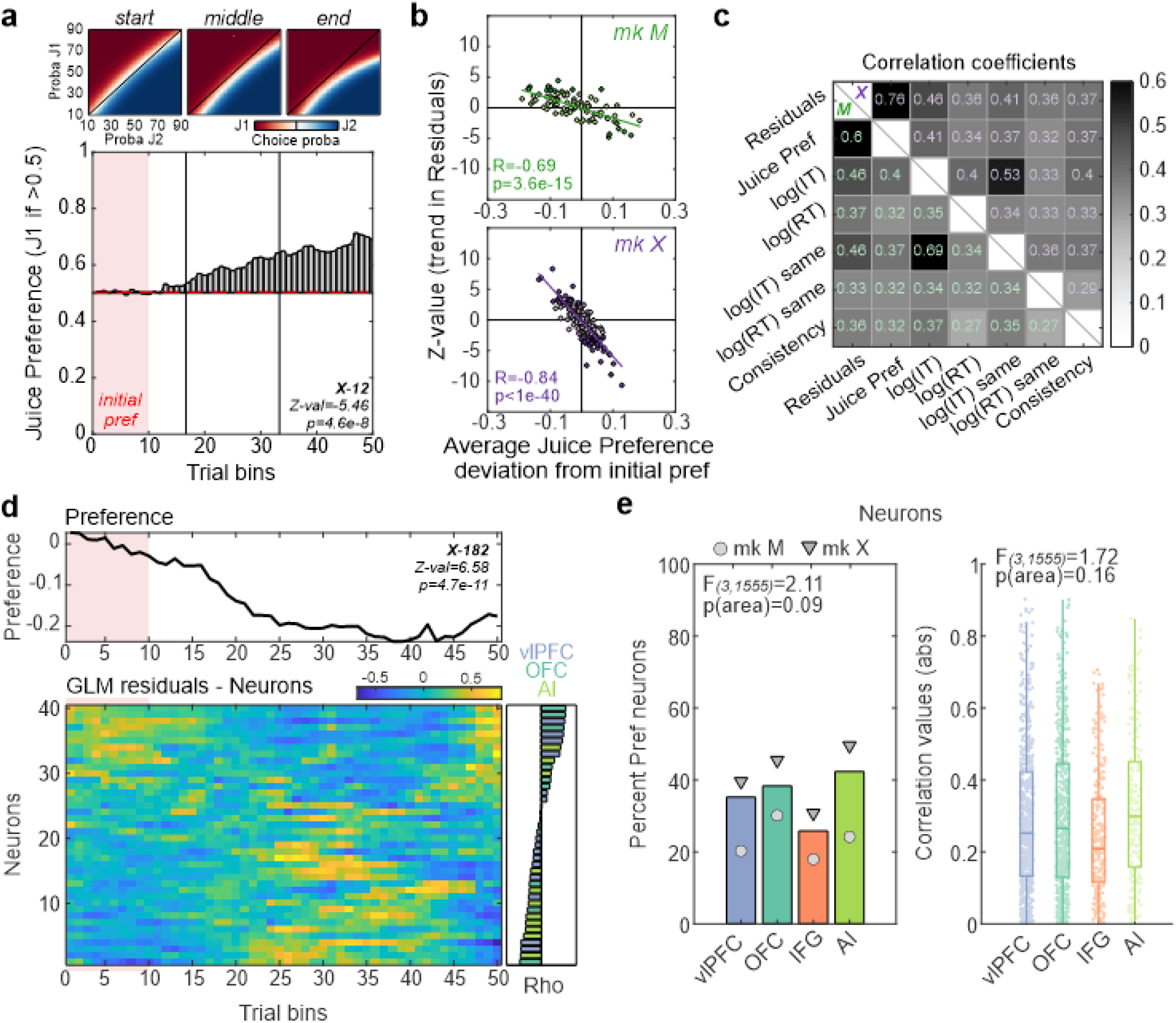
Within-session changes in juice preference and related neural changes. (**a**) Example session from monkey X illustrating change of juice preference over time. Top panels show the modelled choice probabilities for both outcome flavors across session’s terciles, for illustration purposes. Bottom panel shows the juice preference over the course of a session (divided into 50 equally sized bins of trials). Light pink overlay represents the 10 bins used to extract the reference for behavioral/neural measures (e.g., initial preference). (**b**) Across-sessions trend statistics (Z-value) of the behavioral models’ residuals against the average juice preference deviation from the reference period, for monkey M (green, top panel) and X (violet, bottom panel). Each dot represents one session, with the sessions showing a significant trend in residuals highlighted in dark colors (at FDR-corrected p<0.05). (**c**) Average within-session correlation coefficients between all considered behavioral variables (see Methods for variable description), for each monkey (monkey M and X, below and above diagonal respectively). Variables ‘log(IT) same’, ‘log(RT) same’ and ‘Consistency’ were derived from same-flavor trials (e.g., J1 vs J1 trials). Residuals were better explained by the juice preference than any other variables. (**d**) Example session from monkey X showing the time evolution of the preference (top panel) and the residual stimulus-related activity of simultaneously recorded neurons (bottom panel, sorted by correlation strength). We first removed the potential influence of alternative parameters (log(IT) and log(RT)) by using a generalized linear model on neurons’ firing rate and extracting the residuals. Reference period is highlighted in light pink. Bottom right panel shows the correlation coefficient for each neuron, with the color indicating the neuron’s area. (**e**) Left panel shows the percent of neurons across areas showing significant residual firing rate correlation with preference changes (FDR-corrected p<0.05) during the stimulus period. Right panel shows the absolute correlation values for every neuron across areas. The reported statistics were based on mixed-effect logistic (left) or linear (right) regressions.

### The impact of within session changes in juice preference on neural activity

OFC is heavily implicated in updating the value of currently available rewards (Gottfried et al., 2003; Pickens et al., 2003; Izquierdo and Murray, 2004; Rudebeck et al., 2013; Gardner et al., 2017; Howard and Kahnt, 2017; Costa et al., 2023), but how other areas in frontal cortex update the value of specific juices on a moment-to-moment basis is less clear. To establish the impact of juice preference on neural activity we conducted two complementary but separate analyses. The first looked for single neuron activity that correlated with local changes in preference within a session. The second characterized the effect of within-session preference changes on single neuron encoding of outcome flavor or probability, seeking to determine the influence of changing flavor preferences on the specific circuits engaged as subjects made their choices.

To test whether within-session variation in neural activity tracked changes in preference, we first averaged and normalized the activity of each neuron over the same trials used to extract the local preference in sessions with changes in preference (**Fig. 6d** and **Supplementary Fig. 8a**). This gave us a total of 1,559 neurons across all areas on which to conduct our analyses (**Supplementary Table 1**). To account for potential modulation of neural activity with factors related to task execution, we regressed out the influence of log(IT) and log(RT) and used the residual activity for each neuron. Across areas, 35.5% (554/1,559) of neurons showed a significant correlation with within-session preference changes (**Fig. 6e**, left panel, FDR-corrected p<0.05). The proportion of neurons from vlPFC and IFG were the lowest compared to OFC and AI, albeit not significantly different. The strength of the correlations showed large variability across sessions but were consistent between areas (**Fig. 6e**, right panel). Note that analyses accounting for potential drift in our recordings based on waveform change over time did not alter these results (see Methods), and that a similar effect was found at the level of neural populations (**Supplementary Fig. 8b-c**). Thus, within-session effects of preference on neural activity appear to be ubiquitous across the areas recorded.

The prior analysis revealed how preference was tracked at the level of all neurons that we recorded from the frontal lobe. However, these analyses ignore the impact of juice preference on neurons that are likely central to guiding choices; those encoding outcome flavor and probability. The influence of preference on this encoding has the potential to reveal the parts of the frontal cortex that are involved in actively updating the integrated representation of subjective value related to juice flavor.

To assess this, we looked for sessions in which the monkeys’ juice preference exceeded a threshold of 2.5 standard deviations for 10 consecutive trial bins (20% of a session’s length) from an initial 10 bins period. We refer to these task periods as PRE and POST satiety. Such changes in preference were found in 49.5% of all sessions (monkey M=38/97 and monkey X=102/186) where 3,391 neurons were recorded (see **Supplementary Table 1**). Two example neurons recorded in OFC and AI are shown in **Fig. 7a-b**. Both neurons were not only tuned to chosen probability but also showed a decrease in firing rate during the POST period, highlighting the interaction between representations of probability and juice preference.

**Fig. 7.**
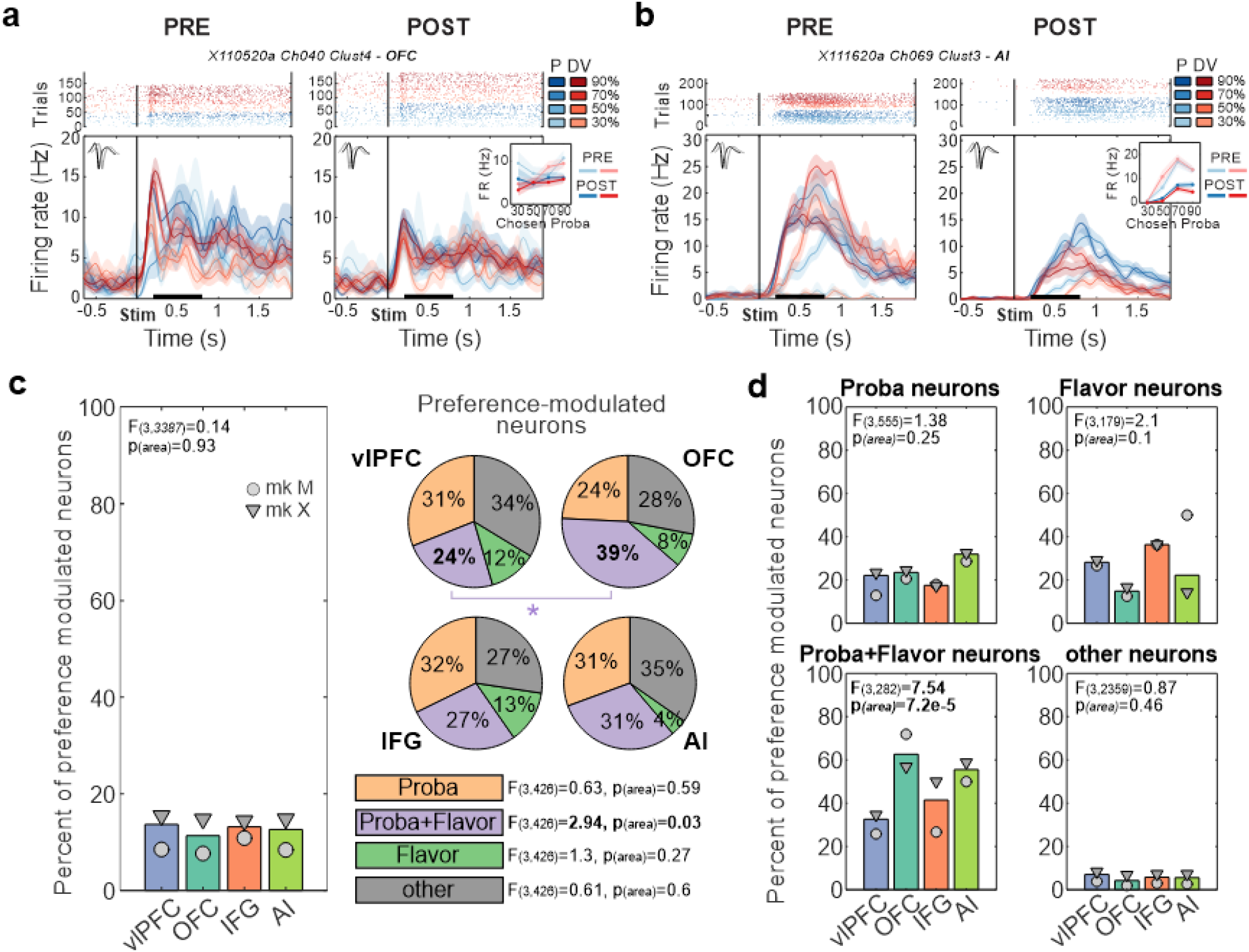
Neural activity related to within-session preference periods. (**a-b**) Raster plots and peri-stimulus time histograms aligned to the stimulus onset for two example neurons recorded in OFC (**a**) and AI (**b**). Left and right panels show the neurons’ responses during the PRE and POST periods, respectively. Colors represent the two possible outcome flavors and the respective chosen outcome probabilities (blues for newly preferred outcome flavor P, red for the devalued flavor DV, and lighter/darker colors for lower/higher probabilities). Insets show the average firing rates across chosen probabilities during the stimulus period (black bar on x-axis) and for all conditions (PRE vs POST periods in light vs dark colors, preferred vs alternative flavor, blue vs red). (**c**) Percent of neurons modulated by preference periods across areas (left panel, at p<0.01) and the proportion of these preference-modulated neurons that were characterized as encoding chosen flavor (green), chosen probability (orange), both (purple) or related to other parameters (grey) for each area (right panel). The star represents a significant difference in proportion between areas, assessed using mixed-effect binomial models (FDR-corrected p<0.05). (**d**) Percent of neurons modulated by preference periods depending on their encoding properties (chosen probability neurons only, chosen flavor neurons only, chosen probability and flavor neurons, or other neurons).

Overall, 12.7% (430/3,391) of all neurons exhibited differences between the two preference periods (p<0.01), with no differences between areas (**Fig. 7c,** left panel). This was greater than chance levels as assessed using permutations and consistent even after accounting for possible waveform changes (see Methods). Within this population of preference-modulated neurons, we found that more neurons were classified as representing both chosen flavor and probability in OFC compared to vlPFC (**Fig. 7c**, right panel; FDR-corrected post-hoc: OFC vs vlPFC, W=2.9, p=0.022). Further analyses revealed that a substantial proportion of preference-modulated neurons showed interactions between preference periods and chosen flavor or probability, most notably in OFC (**Supplementary Fig. 9a**).

Finally, we conducted this analysis in the opposite way; by first classifying neurons as encoding outcome flavor and/or probability and then establishing if their activity was modulated by within-session changes in preferences. Here we found that neurons in all areas were more likely to show preference modulation if they encoded chosen flavor or probability (**Fig. 7d**). The highest proportion of neurons showing such modulation was, however, the small population of neurons representing both chosen flavor and probability in OFC and AI (**Fig. 7d**, FDR-corrected post-hoc: OFC vs vlPFC/IFG, W>2.29, p<0.046; AI vs vlPFC, W=2.37, p=0.046).

In summary, these analyses show that within-session changes in juice preferences impacted the representation of chosen flavor and probability at the level of single neurons. Contrary to what we observed when considering trait-like preferences, neurons exhibiting mixed selectivity for flavor and probability encoding in OFC and AI were the most affected when juice preferences changed (a state-like preference). This finding further bolsters the evidence pointing to a specific role for OFC in encoding the specific moment-to-moment value of the flavor of the predicted outcome (Critchley and Rolls, 1996; Bouret and Richmond, 2010; Howard and Kahnt, 2017; Pastor-Bernier et al., 2021).

## DISCUSSION

By recording across different parts of the frontal cortex and amygdala while monkeys made choices based on the flavor and probability of receiving juice rewards, we tested whether preferences are integrated into a single or separate valuation systems. Our data provide evidence for the latter. Specifically, neurons in OFC more strongly encoded outcome flavor compared to neurons in other areas, whereas neurons in vlPFC and IFG more strongly encoded outcome probability (**Fig. 3**). In amygdala, neurons encoded both aspects reliably, complementing prior lesion works. Our analyses further revealed that the representations of outcome flavor and probability in OFC were the least integrated, as these aspects of the task occupied largely separate activity subspaces (**Fig. 4**). Crucially, frontal cortex representations of outcomes flavor and probability were modulated by monkeys’ preferences for specific juices and these preferences varied at multiple timescales (**Figs. 5-7**). The strength of monkeys’ long-held, trait-like preferences not only correlated with the degree to which outcome flavor was represented in all areas but also differentially influenced outcome probability representations in vlPFC and OFC (**Fig. 5**). Notably, preferences altered the degree to which outcome probability for the least favored juice could be decoded in vlPFC (**Fig. 5d**), whereas it had a non-specific negative impact on encoding of outcome probability in OFC (**Fig. 5c**). Furthermore, local state-dependent preferences within a session had a ubiquitous influence on neurons in all areas (**Figs. 6-7**), with stronger modulations in neurons that encoded flavor and probability within OFC and AI. Thus, by probing how preferences are integrated in neural representations within the frontal cortex, our findings indicate that there are separate valuation systems in frontal cortex.

### (1) Separable subjective valuation systems in frontal cortex for choice

The notion that OFC, primarily Walker’s areas 11, 13 and 14, is a central hub for valuation is based on converging lines of evidence from fMRI, neurophysiology, and lesion studies across species (Gottfried et al., 2003; Izquierdo and Murray, 2004; Padoa-Schioppa and Assad, 2006; O’Neill and Schultz, 2010; Stalnaker et al., 2014; Rich and Wallis, 2016; Gardner et al., 2017; Howard and Kahnt, 2017; Pastor-Bernier et al., 2019; Setogawa et al., 2019; Costa and Averbeck, 2020; Kuwabara et al., 2020). Because of this, there has been an intense focus on determining the mechanisms of valuation in this one part of frontal cortex. Another reason for an overemphasis on the OFC is the fact that this part of frontal cortex has clear homologues across rodents and primates whereas vlPFC does not (Preuss and Wise, 2022). As a result, less attention has been paid to vlPFC, which is anatomically (Carmichael and Price, 1994) and functionally distinct from OFC (Murray and Rudebeck, 2018; Monosov and Rushworth, 2022) and how the former might signal subjective value.

Here we report that neurons in vlPFC are more likely and more strongly encoding the probability of receiving a reward compared to OFC (**Fig. 3**). Of course, this does not necessarily mean that this area is encoding subjective valuations. However, the finding that stable trait-like preferences scale with the ability to decode the probability of the least preferred options only in vlPFC (**Fig. 5**) does indicate that preferences are being integrated to derive the subjective value of the available options. Notably, vlPFC-linked valuations appear to be referenced to the probability of receiving a reward. Whether other parts of frontal cortex encode valuations in distinct reference frames, and integrate preferences into these representations, is an open question, but several findings indicate that they do (Rudebeck et al., 2008; Kennerley and Wallis, 2009; Stoll et al., 2016; Hunt et al., 2018).

In contrast, while preferences scaled with representations of outcome flavor encoding across all recorded areas (**Fig. 5**), it was primarily activity in OFC, and to a lesser degree AI, that closely tracked the within-session changes in how monkeys valued each of the juices (**Figs. 6**-**7**). This finding indicates that representations in OFC and possibly AI are more closely linked to the outcome flavor associated with the available juices and its updated, subjective valuation at any given time. Such a role for these two parts of ventral frontal cortex in representing outcome-flavor linked valuations would fit with the known anatomical connections of these areas, with OFC and AI receiving strong olfactory, gustatory and visceral sensory information (Carmichael and Price, 1995a, 1995b, 1996). It would also help to explain why many prior studies placed OFC as central to subjective valuations: the majority of these manipulated animals’ preferences for specific outcomes such as food rewards or juices by devaluing them through satiety or taste aversion (Nakano et al., 1984; Critchley and Rolls, 1996; Pritchard et al., 2008) or had subjects’ trade-off between different amounts of distinct fruit juices (Padoa-Schioppa and Assad, 2008; Padoa-Schioppa, 2009; Pastor-Bernier et al., 2019).

While we found distinction between OFC and vlPFC in the representation of flavor and probability, it was notable that neurons in amygdala robustly represented both of these aspects of the task (**Fig. 3**). Amygdala, especially the basal nucleus, projects to OFC and vlPFC (Carmichael and Price, 1995a; Ghashghaei et al., 2007) and a recent analysis shows that distinct populations of amygdala neurons send projections to these two parts of the ventral frontal cortex (Zeisler et al., 2023). This potentially indicates that there are anatomically separable circuits between the amygdala and frontal cortex; OFC-amygdala interaction is required for updating the current value of distinct flavors (Baxter et al., 2000) whereas vlPFC-amygdala interaction could be required for updating estimates of reward probability (Chau et al., 2015).

The differences in value representations in OFC and vlPFC provide insight into how frontal cortex-linked valuations guide choices. We found that representations in vlPFC showed more complete integration of flavor and probability compared to OFC (**Fig. 5**). One interpretation of this observation is that vlPFC functions at a higher hierarchical level than OFC in terms of representations of subjective value. Such representations could then drive behavior by relaying this information to prefrontal and premotor areas more closely associated with guiding actions (Yeterian et al., 2012; Saleem et al., 2014; Trambaiolli et al., 2022), such as IFG. Indeed, encoding within IFG diverged from the other areas that we studied as it more strongly encoding the response direction on each trial (**Supplementary Fig. 2)**. Notably, probability information was also reliably signaled in this area, but these representations were not affected by preferences (**Fig. 5c-d**). As such, signals in IFG appear to be less related to the subjective value of the options being chosen and instead may reflect attentional processes or movement planning. If representations in IFG are more closely linked to overt attention this could potentially help to explain why flavor representations in IFG strengthened when subjects exhibited preferences for one juice over another (**Fig. 5b**).

### (2) Everything is not everywhere and even when it is, it is not represented similarly

Neurophysiological studies of ventral frontal cortex have long been constrained to acquiring simultaneous recording from a very limited set of areas. Consequently, researchers have predominantly focused on discerning how activity within a single area is correlated with different aspects of a task. Because of this it has been hard to establish a complete picture of where information, such as subjective value, might be the best represented. Indeed, some researchers have argued for distributed value representations in the brain (Hunt and Hayden, 2017). The approach we took here meant that we were able to 1) study multiple areas across frontal cortex and amygdala simultaneously and 2) record enough neurons in each area to conduct neuronal analyses on single sessions. Overall, we found 20-70% of neurons significantly correlating with any given task parameter of interest in our subjects. Such a result would appear to support the idea that reward attributes are signaled in a distributed manner; not only could correlates of the various task parameters be found across the amygdala and multiple frontal areas but individual neurons within these areas were also more likely to represent multiple parameters (**Fig. 4**). Such mixed selectivity, the co-representation of multiple variables in the activity of single neurons, was a prominent feature in our data, supporting previous observations within frontal and subcortical areas (Mante et al., 2013; Rigotti et al., 2013; Saez et al., 2015; Putnam and Gothard, 2019).

However, the observation that a large number of neurons in an area had significant modulation for a valuation-related variable did not necessarily predict the strength of these representations. This was especially true for representations of outcome flavors as similar numbers of neurons in OFC, vlPFC, IFG, and amygdala were classified as encoding outcome flavor, but activity in OFC neurons was more strongly tuned to outcome flavor and decoding performance in this area was higher compared to others (**Fig. 3**). In addition, when we looked at how outcome flavor and probability were represented in each area, there were clear differences in how separable the activity subspaces were across areas, with representations of these two task parameters being most separable in OFC (**Fig. 4**). Finally, when we looked at how preferences influence encoding, there were marked differences between areas; trait-like preferences impacted encoding in a circumscribed set of areas whereas within-session (state-dependent) changes in preference had their greatest impact in OFC. Taken together, our findings point to specialized representations of value in different parts of frontal cortex and argue against distributed coding accounts.

### (3) Summary and conclusion

Our data indicate that there are distinct representations of subjective value in OFC and vlPFC, but why there is such a separation is an open question. One reason may relate to the way that some nonhuman primates, and specifically macaques, forage for food (Murray et al., 2020; Monosov and Rushworth, 2022; Rudebeck and Izquierdo, 2022). Macaques forage over large day ranges encompassing many thousands of hectares. They use vision to seek out locations where high-calorie foods might be obtained. However, it is only when a food can be closely inspected and consumed that its qualities can be discerned, including olfactory and tactile sensations that indicate features such as ripeness, which is strongly associated with objective value (i.e., caloric content). Consequently, vlPFC-linked probability representations may be more engaged during foraging choices made at a distance, which necessarily entail uncertainty about outcomes, whereas foraging in close proximity to food items or flavored fluids requires OFC-linked systems to predict the outcome-specific qualities of a given choice. This distinction would fit with known differences in the anatomical connections of these areas; OFC receives inputs from olfactory and gustatory areas, whereas vlPFC receives dense input from visual areas in the inferior temporal cortex (Carmichael and Price, 1995a, 1995b). Both of these systems would need to integrate preference information to optimally guide choices to desirable foods or other items of interest.

## ACKNOWLEDGEMENTS

This work was supported by a National Institute of Mental Health BRAINS award to PHR (R01 MH110822), a young investigator grant from the Brain and Behavior Foundation (NARSAD) to PHR, a Philippe Foundation award to FMS, and seed funds from the Icahn School of Medicine at Mount Sinai to PHR.

## METHODS

### Subjects

Two adult male rhesus macaques (*Macaca mulatta*), monkeys M and X, served as subjects. They were 8 and 5.5 years old, and weighed 11.9 and 7.9 kg, respectively, at the start of the neurophysiological recordings. Animals were grouped housed, kept on a 12-h light dark cycle and had access to food 24 hours a day. Throughout training and testing each monkey’s access to water was controlled for 5 days per week. All procedures were reviewed and approved by the Icahn School of Medicine Animal Care and Use Committee.

### Apparatus

Each monkey was habituated to sit in a custom primate chair with their head restrained, situated 56 cm from a 19-inch monitor screen. Choices were indicated by gaze orientation. Eye movement and pupil size were monitored and acquired at 90 frames per second using an infrared oculometer (PC-60, Arrington Research, Scottsdale, AZ). Juice rewards were delivered to the monkey’s mouth using a custom-made air-pressured juice dispenser system (Mitz, 2005). Trial events, reward delivery and timings were controlled MonkeyLogic behavioral control system (https://monkeylogic.nimh.nih.gov), running in Matlab (version 2014b, The MathWorks Inc.). Raw electrophysiological activity was recorded using an Omniplex data acquisition system (Plexon, Dallas, TX) and sampled at a 40kHz resolution. Spikes from putative single neurons were automatically clustered offline using the MountainSort plugin of MountainLab (Chung et al., 2017) and later curated manually based on the principal component analysis, interspike interval distributions, visually differentiated waveforms, and objective cluster measures (Isolation probability > 0.75, Noise overlap probability < 0.2, Peak signal to noise ratio > 0.5 sd, Firing Rate > 0.05 Hz). Most neurons in our dataset were highly isolated (**Supplementary Fig. 1**, median; Isolation probability = 0.986, Noise overlap probability = 0.013, Peak signal to noise ratio = 5.73 sd, Peak noise = 1.67 sd, Firing Rate = 2.28 Hz). Behavioral and neurophysiological analysis were performed offline using custom Matlab scripts.

### Behavioral tasks

Monkeys were trained to perform 3 closely related tasks during each session, including a single option, Instrumental (**Fig. 1a**) and Dynamic probabilistic tasks. The stimuli used were common to the different tasks and contained two features: an external-colored rectangle indicating which outcome flavor (out of 2 possible juices per session) monkeys could earn and a second central rectangle, more or less filled, indicating the probability at which this particular outcome flavor would be delivered at the end of the trial. During a given session and across tasks, monkeys were faced with options containing one of two possible colors (randomly selected from a set of 9 colors) associated with two different juice flavors (randomly picked from a set of 5, which included apple, cranberry, grape, pineapple and orange juices, diluted in 50% water). Probabilities used were from 10% to 90% (by steps of 20% for monkey M and 10% for monkey X).

Each session started with a defined number of trials of the single option task (monkey M, n=115; monkey X, n=300 trials) and was followed by blocks of 4 Instrumental trials intermixed with 1 Dynamic trial (i.e 4Ins-1Dyn-4Ins-1Dyn-etc.). Behavioral analyses were restricted to sessions with a minimum of 100 different juices Instrumental trials (monkey M, 103 sessions, min/median/max=104/184/292 trials; monkey X, 186 sessions, min/median/max=239/802/1336 trials). Performance and neurophysiology during the single option and dynamic tasks were not considered in the current analyses (except for the previous chosen flavor parameter, see Behavioral analyses).

#### Instrumental task

In this task, monkeys were operantly conditioned to choose between two options presented simultaneously, one on the left side of the screen and the other on the right side. Each option was associated with particular outcome flavor and a probability of receiving it, and a visual stimulus indicated the options, as described above.

Monkeys initiated a trial by fixating a central fixation cross for 0.7 to 1.3s (steps of 0.3s). The two options, each composed of an outcome flavor and probability, were then displayed for 0.4 to 0.8s (steps of 0.2s), pseudorandomly. The stimulus was then turned off for 0.2s and two response boxes appeared on both sides of the previously shown options (3 possible locations equidistant to the options’ locations; bottom left/right, center left/right, top left/right). Monkeys had to fixate the response box on the side of the desired option to make their choice. Fixation on one of the response boxes had to be initiated within 8s. Monkeys needed to acquire and maintain fixation on the selected response box for a minimum of 0.25s in order to register a response, at which time the other one would disappear. The dwell time enabled subjects to change their choice within a trial. After response registration, fixation was required for an additional 0.6 to 1.2s (steps of 0.3s). At that point, the response boxes would disappear for 0.3 to 0.7s (steps of 0.2s). At the time of the feedback, both options were presented again at the same locations, with the selected one initially flashing (5 times 0.1s ON followed by 0.1s OFF, total time of 0.5s, indicated by 5x in **Fig. 1a**), before staying on the screen for the duration of the reward (if delivered) and an additional 0.5s. In rewarded trials, monkeys received 2-3 pulses of 0.03-0.06s of fluid (separated by 0.1s each, 0.25-0.36ml total reward per trial of the outcome flavor (and at the probability) indicated by the selected option). Nonrewarded trials were matched in time to trials that included reward delivery. Finally, rewarded trials were followed by a 2s intertrial interval, and unrewarded trials were followed by 3.5-4s. If monkeys failed to maintain fixation when required, a large red circle was presented at the center of the screen for 1s, followed by a longer intertrial interval (4-6s for monkey M, 3-4s for monkey X). Failure to initiate a trial by looking at the fixation star within 6s of its appearance resulted in the same red circle and intertrial interval.

The two options were associated with different outcome flavors (juice 1 vs. 2) in half of the trials for monkey M and in 3 out of 4 trials for monkey X. In these trials, the probability was always different for the two outcomes in monkey M but could be either different or similar in monkey X (e.g., juice 1 at 70% vs. juice 2 at 70%). In the remaining trials (1/2 for monkey M and 1/4 for monkey X), the two options were associated with the same outcome flavor (e.g., juice 1 vs. juice 1) but with different probabilities.

#### Single option and Dynamic tasks

Monkeys were also asked to complete a single option task at the start of the experiment and either a Dynamic single option task (monkey X) or an Instrumental Dynamic task (monkey M). The trial structure and event timings were similar to the Instrumental task except as reported below.

In the single option task, monkeys were shown a single option (either on the left or right, pseudorandomly selected) which was associated with a particular outcome flavor and probability.

In the Dynamic single option version, monkey X was shown sequentially two options associated with different probabilities but on the same side of the screen (option 1 for 0.4 to 0.8s, followed by a blank screen for 0.2s, and then option 2 for 0.4 to 0.8s also followed by a blank screen for 0.2s; total time = 1.2 to 2s, steps of 0.4s). Both options were always associated with the same outcome flavor. After maintaining fixation on the response box, the average probability associated with the two successive options was given as feedback (e.g., if option 1 was associated with 40% reward and option 2 with 80%, the option during feedback was associated with 60%), which defined the probability of getting a reward at the end of the trial. Only 4 probability sequences were used here (0% and 40%; 0% and 100%; 40% and 80%; 80% and 100%), resulting in 4 different final probabilities (20%; 50%; 60% and 90%, respectively).

In the Instrumental Dynamic version, monkey M was shown sequentially three pairs of options associated with different probabilities. Specifically, a pair of options was first presented simultaneously on both side of the screen (as in the Instrumental task; pair 1 for 0.4 to 0.8s, followed by a blank screen for 0.2s), followed by pair 2 and pair 3 (same durations; total time = 1.8 to 3s, steps of 0.4s). The monkey was then free to fixate one of the two response boxes. At the time of feedback, the average probability for both options was presented simultaneously, with the selected option flashing, like in the Instrumental task. In this task, only a subset of probabilities was used (30-70-80 vs. 90-90-60; 10-60-80 vs. 90-90-60; 10-50-90 vs. 90-90-30; 10-10-70 vs. 50-90-10; 10-10-70 vs. 50-80-20; 10-50-60 vs. 90-30-90; 30-30-90 vs. 80-90-70; 30-30-90 vs. 70-80-60; 10-20-60 vs. 70-70-10), resulting in 5 different final probabilities (30% vs. 50%; 40% vs. 70%; 50% vs. 80%; 50% vs. 70%; 60% vs. 80%). The two options within a pair were associated with the same outcome flavor in 2 out of 3 trials. In the remaining trials (1/3), monkeys were faced with different flavors on both sides. In both cases however, the 3 sequentially presented options on a given side were always associated with 1 flavor (i.e., A-A-A vs. B-B-B, A-A-A vs. A-A-A or B-B-B vs. B-B-B; where A and B are the 2 possible outcome flavors).

#### Selective satiation task

Monkey X completed a selective satiation task (n=6 sessions, **Supplementary Fig. 7a**). Here, the animal first performed 100 trials of the single option task, followed by 550-600 trials of the Instrumental task. After that, a large 200-250ml bolus of one of the two juices available during that session was given to the animal within a 15min period. The size of this bolus was equivalent to ∼40-50% of this animal’s daily liquid consumption. Following this bolus, the animal was asked to complete an additional 600-650 trials of the Instrumental task.

### Surgical procedures and neural recordings

All surgical procedures were conducted under full anesthesia in a dedicated operating room under aseptic conditions. Anesthesia was induced with ketamine hydrochloride (10 mg/kg, i.m.) and maintained with isoflurane (1.0-3.0%, to effect). The animal received isotonic fluids via an intravenous drip. We continuously monitored the animal’s heart rate, respiration rate, blood pressure, expired CO2 and body temperature. The animals were treated with dexamethasone sodium phosphate (0.4 mg/kg, i.m.) and cefazolin antibiotic (15 mg/kg, i.m.) for one day before and for one week after surgery. At the conclusion of surgery and for two additional days, the animals received ketoprofen analgesic (10-15 mg/kg, i.m.); ibuprofen (100 mg) was administered for five additional days.

Monkeys were first implanted with a titanium head restraint device, fixed to the cranium with 14 titanium screws. Based on T1-weighted magnetic resonance images of the monkeys’ brain, we then placed a form-fitted plastic recording chamber over the exposed cranium of the right frontal lobe of both monkeys, attached to the cranium using both titanium screws and dental acrylic (C&B Metabond, Parkell Inc, Edgewood, NY; Ortho-JET BCA, Lang Dental MFG Co., Wheeling, IL). We acquired another MRI where the chamber was filled with gadolinium saline (Magnevist, Bayer, 1:1000 dilution) and mounted with external fiducials. This procedure allowed us to adjust the path of each electrode based on the exact placement of the chamber. In a following surgery under general anesthesia, we performed a craniotomy inside the chamber. After confirming the absence of bacterial growth inside the chamber (anaerobic and aerobic cultures, IDEXX BioAnalytics), we performed a final surgery consisting of first puncturing the dura where the electrodes would penetrate the brain before sealing the chamber by placing a 157-channel semi-chronic microdrive system (Gray Matter Research, Bozeman, MT; see **Fig. 2a**) containing glass-coated electrodes (1-2MΩ at 1kHz; Alpha Omega Engineering, Nazareth, Israel). Electrodes were immediately lowered beyond the dura to minimize electrode tip damage. Further details on the procedure and microdrive can be found at https://www.graymatter-research.com/documentation-manuals (Dotson et al., 2017). Only electrodes with impedance >0.2MΩ at 1kHz were further lowered and recorded (monkey M: n=116/155, monkey X: n=105/151, the difference in the denominators reflects electrodes that were not lowered given possible risks associated with their trajectories).

Due to the world-wide COVID-19 pandemic, recordings of monkey X were paused for a few months. Given the relatively low number of electrodes and the time lost, we decided to replace the microdrive. Before explanting, we acquired CT images with 7 electrodes left in place. The first microdrive was then removed and we collected an additional structural MRI to assess possible brain displacement and map our electrode locations. Following a craniotomy and additional duratomy, an identical 157-channel semi-chronic microdrive system was placed and electrodes were immediately lowered (n=137/151).

Recording locations were confirmed in both monkeys using several approaches. First, we recorded the cumulative depth of each electrode while slowly lowering them on a regular basis, tracking changes in background noise and electrophysiological activity suggesting white/gray matter transitions. We also acquired CT images at different time points, which were then co-registered to both the post-operative MRI and an additional MRI performed at the conclusion of the experiment. Finally, we captured and digitalized block-face pictures before cutting every brain section which were later stained, showing clear marks of the electrodes’ track. Combined with microscope observations of histological stained sections (see below for more details) and using Free-D software (Andrey and Maurin, 2005), this allowed us to reconstruct the anatomical location of every electrode.

### Tissue preparation and immunohistochemistry

At the conclusion of the neurophysiology recordings, the monkeys were deeply anesthetized (ketamine, 10 mg/kg and sodium pentobarbital, 100 mg/kg) and transcardially perfused with 4% formaldehyde (EMS, Hatfield, PA) in Phosphate-buffered saline (PBS; Fisher Scientific, Fair Lawn, NJ). The brain was extracted and post-fixed in buffered formaldehyde (4% by weight) for 24-48h. The brain was then placed in increasing concentration of sucrose solutions (10%, 20%, and 30% sucrose in PBS), waiting for it to sink at every step, before being shipped for sectioning and staining.

The following cryopreservation, sectioning, mounting, tissue staining, and immunohistology services were all performed by FD NeuroTechnologies, Inc (Columbia, MD). Specifically, the recorded brain hemispheres were cryoprotected in FD tissue cryoprotection solution^TM^ for 10 days before being rapidly frozen in isopentane pre-cooled to -70°C with dry ice. Serial sections (50 μm) were then cut coronally through the right hemispheres, approximately from +48 mm to +15mm interaural (Saleem and Logothetis, 2012) with a sliding microtome. Every 1st, 2nd, 3rd, and 4th sections of each series of 4 sections (interval: 200 μm) were collected and stained separately. The first series was stained with FD cresyl violet solution^TM^ (Nissl staining, 1 section every 200 μm). The second and third series were processed for Calbindin (using mouse monoclonal anti-Calbindin-D-28K antibodies, 1:1000 dilution; Millipore Sigma, St. Louis, MO) and SMI-32 (using mouse Purified anti-Neurofilament H nonphosphorylated antibodies, 1:12000 dilution; Biolegend, San Diego, CA) immunohistochemistry (note that only sections adjacent to every 3rd Nissl-stained sections were processed for calbindin or SMI-32, meaning 1 section every 600 μm). Specifically, after inactivating the endogenous peroxidase activity with hydrogen peroxidase, sections were incubated separately with avidin and biotin solutions (Vector Lab, Burlingame, CA) for blocking nonspecific binding of endogenous biotin, biotin-binding protein and lectins. Subsequently, sections were incubated, free-floating, in 0.01 M PBS containing Triton X-100, blocking serum, and the primary antibody for 67 hours at 4°C. The immunoreaction product was then visualized according to the avidin-biotin complex method (Hsu et al., 1981) using the Vectastin elite ABC kit (Vector Lab., Burlingame, CA) and 3’,3’-diaminobenzidine as a chromogen. After thorough washes, all sections were mounted on microscope slides, dehydrated in ethanol, cleared in xylene, and coverslipped in Permount (Fisher Scientific, Fair Lawn, NJ).

### Defining neuroanatomical boundaries

Neuroanatomical boundaries were drawn and measured from histological sections, and we referred below to the various considered subdivisions and landmarks (Saleem and Logothetis, 2012). Ventrolateral prefrontal cortex (vlPFC) recordings included all subdivisions of area 12 (rostral, medial, lateral and orbital) as defined by Carmichael and Price (1994). Orbitofrontal cortex (OFC) recordings were mainly limited to the portion between the medial and lateral orbital sulcus, covering most of area 11 and 13 (with the exception of 13b). Inferior frontal gyrus (IFG) consisted of area 44 and 45 (dorsal to the lower limb of the arcuate sulcus). Agranular insular cortex (AI) included recordings from the intermediate, lateral, medial and posteromedial subdivisions. Finally, amygdala recordings were mostly focused on the basolateral portion but included surrounding subdivisions. Neurons recorded more than 0.2mm outside of the boundaries defined from histology sections were discarded.

### Behavioral analyses

#### Choice patterns during Instrumental task

Behavior during the Instrumental task was analyzed using logistic regressions (function *fitglm* in Matlab). Specifically, the odds of choosing one of the 2 flavors in trials where the two options were associated with different flavors were fitted using the following model:

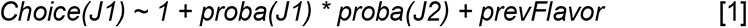

where *proba(J1)* and *proba(J2)* are the probability (linear variable) associated with both outcome flavors and *prevFlavor* (categorical) the last chosen outcome flavor in the preceding trial. Note that the last chosen outcome flavor was independent of which task or condition was presented or whether the animal received it or not. In case no options were presented on the previous trial (break of fixation or no trial initiation), the previous chosen flavor was used. All variables in the model were normalized between -1 and 1. We extracted the significance and estimates for each variable, the predicted choice probabilities (with *proba(J1)* and *proba(J2)* from 10% to 90% with 1% steps and *prevFlavor* set to -1), as well as the models’ residuals for every session. Sessions where the model did not converge were excluded from further analyses (monkey M, n=97/103; monkey X, n=186/186 sessions).

For each session, monkeys’ juice preference was extracted by computing the proportion of choices where the probability of choosing flavor J1 was higher than flavor J2, using the predicted choice probabilities from our behavioral model.

To ensure outcome preference were not idiosyncratic but goal-directed, we assessed whether monkeys’ outcome juice preference across sessions followed transitivity using all possible triplet of juice: if monkeys preferred flavor A over B and B over C, then they should prefer A over C. Given that monkeys were faced with only 2 outcome flavors on a given session, we matched all possible sessions where monkeys were presented with flavor A-B, B-C and A-C. We then extracted the average of the product of the odds of choosing A over B (in a given session) and B over C (in another session) with the average odds of choosing A over C (in a third session). We tested for difference across triplet of juice flavors and monkeys using two-sided sign test.

#### Within-session behavioral changes

Change of preference occurring within a session were first assessed using a Mann-Kendall trend test of the models’ residuals (at FDR-corrected p<0.05), an indicator that the models’ fit was not appropriate across trials. To specifically quantify monkeys’ outcome juice preference over time, we fitted their choices over 50 sliding windows (with a window size of 1/5^th^ of the total number of trials performed) using the same model [1]. When models converged, we were able to extract the local juice preference in a similar manner than for the overall juice preference described above. Finally, we extracted the average deviation of the local juice preference across the whole session, after referencing it to the monkey’s initial preference (using the average of the first 10 sliding windows). This provided an index of their overall change of preference for each session (**Fig. 6c**).

We ensured that changes of preference were independent from changes in motivation or inattention using several approaches. First, we extracted the log-transformed average initiation time (log(IT); time between the fixation cross appearance and the start of fixation) and log-transformed average reaction time (log(RT); time between the response boxes onset and first fixation) across the same 50 sliding windows. These time series were further min-max normalized (between -1 to 1). We also used trials where both options presented were associated with the same outcome flavor (e.g., J1 vs. J1). Specifically, we computed the proportion of choices where monkeys selected the option with the highest probability for each outcome flavor across the same 50 sliding windows, a measure of monkeys’ flavor-specific choice consistencies. We also extracted the initiation and reaction times across these same-flavor trials (log(IT) same and log(RT) same in **Fig. 6c**). These parameters were correlated against each other, and the average correlation across sessions was reported (**Fig. 6c**).

Finally, we also extracted periods when outcome juice preference diverged from the initial preference. For this, we extracted the trials when local preference exceeded a threshold of 2.5 standard deviations for 10 consecutive windows (POST period) when compared to the first 10 sliding windows (PRE period). We ignored the individual trials that were in both periods, which could be the case given the sliding window approach.

#### Choice patterns following selective satiation

We assessed how changes of preference might be modulated in similar ways following selective satiation in monkey X (**Supplementary Fig. 7**). Local juice preference before and after the juice bolus was assessed using the fitted choices over 50 sliding windows in each period, as previously described (model [1]). Initiation and reaction times were also computed for different outcome flavor trials across the same windows. Similarly, Mann-Kendall trend tests were used to assess the existence of trends in the various variables, both before and after the bolus. To assess the existence of a trend deviation before and after bolus, independently of its direction (e.g. toward the juice given as a bolus or away from it), the sign of the preference-related z-values for the period before bolus were forced to be positive (e.g. if the z-values pre/post bolus were -2.5/+5.3, these values were inverted to +2.5/-5.3). Z-values from the trend tests were then compared using Wilcoxon signed-rank tests.

Finally, we extracted the choice probability before and after bolus (and the difference between these) depending on the difference in probability of the two options presented for every different outcome flavor trials. This allowed us to derive the integral of the difference in choice probability across all probability difference, providing an estimate of the relative bias toward or against the given juice following the bolus.

### Electrophysiological data processing

#### Pre-processing

We focused our analyses on the instrumental task, specifically the response of neurons during trials where different outcome flavors were offered. Neurons were included based on the quality of isolation only, with no response pattern or firing rate restrictions. Spiking activity for each trial was first smoothed using a 50ms Gaussian kernel before being averaged over 20ms bins. Neurons’ firing rates were aligned to multiple events across trials (central fixation, stimulus onset, response fixation, feedback and reward onsets). Most of the analyses reported here were related to the stimulus-related activity, notably a period between 200-800ms following the stimulus onset. Results are provided for each monkey as well as combined.

#### ANOVA on single neuron responses

For this analysis, we analyzed the responses of neurons recorded during sessions lasting at least 150 trials. After applying a min-max normalization of the binned firing rate of each neuron, we used four-way ANOVAs to explain the variance in firing rate from each neuron and at every time bin associated with the following factors of interest: Chosen Flavor (2 levels, categorical), Chosen Probability (4 levels, 30-50-70-90%, linear), Unchosen Probability (4 levels, 10-30-50-70%, linear) and Chosen Side (2 levels, categorical). No interactions were included in the models. Neurons were considered as significantly encoding a task factor if they discriminated that factor for 3 consecutive bins (covering a time period of 100ms) with a threshold of p<0.01. This led to a chance level of ∼5% of significant neurons for any given factor when considering the period before the stimulus onset (100-700ms following central fixation) (black line on **Fig. 3a, e**). The onset of significant encoding of a given factor was also extracted for each neuron, considering only the stimulus period (0 to 1000ms).

We computed the effect size, omega-squared, for each factor included in the ANOVA model and using the following formula:

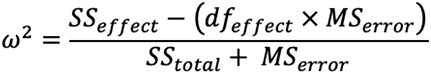

where *SS_effect_*and *SS_total_* are the sum of squares of the considered factor and of all factors (respectively), *df_effect_* represents the degree of freedom associated with the factor, and *MS_error_* the mean squared error. Omega-squared therefore represents an estimate of the amount of variance accounted by the explanatory factor.

Finally, we assessed mixed selectivity across neuronal populations by calculating the proportion of chosen flavor neurons that were also considered chosen probability, compared to the proportion of non-chosen flavor neurons being chosen probability. This ratio was compared with 1000 permutations, where we shuffled the significant/non-significant labels associated with chosen probability across neurons. The reciprocal comparison (i.e. that chosen probability neurons were more likely or not to also be tuned to chosen flavor) led to similar results and was not reported.

#### Population decoding

Population decoding was performed for chosen flavor, chosen and unchosen probability, the chosen probability of flavor J1 and J2 separately and chosen side. We only considered sessions with at least 5 neurons simultaneously recorded in the considered area. In all cases, the activity of all neurons was first averaged around the stimulus period (200-800ms), before extracting a subset of similar number of trials across each possible category (defined as the lowest number of trials across categories, with a minimum of 5). Given that monkeys rarely chose the lowest probability (or rarely avoided the highest), and to ensure the decoding analyses are well-powered, we only considered 4 chosen probabilities (30, 50, 70, 90%) and unchosen probabilities (10, 30, 50, 70%) when decoding these parameters. The trial selection procedure and the following steps were performed 200 times. Dimensionality was reduced by using PCA, and the decoding was then performed by applying linear discriminant analysis (LDA) on the activity captured in the first 4 components using 10-fold cross-validation. The decoding performance on the testing sets were then averaged across the 10-fold cross-validations, before being averaged across the 200 trial-selection permutations. Additionally, we assessed the significance of decoding performance by permuting the trial labels for each trial-selection permutation (resulting in 200 random permutations). Decoding was considered significant if less than 10 random permutation decoding performance were greater than the true average performance (one-tailed, p<0.05).

When investigating the relationship between neural representations and between-session preferences, we correlated the performance of these classifiers with the preference level (or its absolute value when appropriate).

#### Probability and Flavor subspaces

Here, we first applied a min-max normalization of the binned firing rate of each neuron across all trials and time window of interest (from fixation onset to reward offset). To extract the chosen flavor and probability neural subspaces, neurons’ firing rate following stimulus onset (200-800ms) were averaged and centered across a subset of trials (i.e., the training set of a 10-Fold cross-validation procedure). As for the decoding analyses, we included only trials with 4 chosen probabilities and both flavors. The average population activity was used to train a PCA before extracting the angle in N-dimensional space (N being the number of neurons considered) between the two subspaces (function *subspace* in Matlab). We report the average angle (between 0 and 90 degrees) across a 10-fold cross-validation procedure.

For the simulated example in **Fig. 4c**, only 2 neurons were considered, with their firing rate the sum of activity associated with the different probabilities (weights = 5 vs 2), flavors (weights = 1 vs 5) and additional noise, across 10 repetitions of each condition (2 flavors and 4 probabilities). The following steps were similar.

#### Within-session preference changes modulation of single unit and population responses

Our first analysis was limited to recordings made during sessions with significant trend in residual, as previously described (see **Supplementary Table 1** for included numbers, **Fig. 6** and **Supplementary Fig. 8**). Here, we averaged and min-max normalized the stimulus-related activity of each neuron over the same trials used to extract the local preference (see above, 50 sliding window model). Neurons with no average activity across the considered bins were included but not normalized. We then regressed out the influence of execution metrics by applying a generalized linear model (GLM) to explain each neuron’s average activity using factors log(IT) and log(RT). Both factors were mean-centered (using the mean of the initial state, the first 10 sliding windows). The residual activity of each neuron was finally used and Pearson-correlated with the local preference, also mean-centered. Neurons were considered significantly correlated with local preference using a threshold of p<0.05 after FDR-correction.

We also assessed the correlation between changes in preference and population responses by applying PCA to the residual activity of simultaneously recorded neurons (minimum of 5 neurons) for each area. We then derived the population distance over trial bins by computing the Euclidean distance (in 3-dimensional space) of the population neural activity at every window compared to its initial state, the average activity during the first 10 sliding windows. As for single neuron activity, we correlated the population distance with the local preference and used a threshold at p<0.05 FDR-corrected.

Lastly, we ran a complementary analysis, limited to recordings made during sessions with significant deviation of local preference (i.e., PRE vs POST periods, see **Supplementary Table 1** and **Fig. 7**). We first extracted the stimulus-related activity of each neuron during the trials contained within both PRE and POST periods. We then performed 3-way ANOVAs to explain the variance in firing rate of each neuron with the chosen flavor, the chosen probability, and the preference periods (2 categorical levels, PRE vs POST) as predictors, as well as their interactions. In parallel, we ran 100 permutations where we shuffled the firing rate across trials for each neuron, keeping the relationship between the different predictors intact. Neurons were considered significantly tuned to a parameter if they discriminated that parameter at a threshold of p<0.01 in the main analysis. Such threshold led to an average of 3.5% of neurons (min-max, 2.7%-4.3%) from all areas to be considered (wrongly) significant for preference periods (main effect or interactions) across the permutation sets.

#### Waveform stability control

To rule out the possibility that changes in firing rate were related with changes in spiking isolation/detection, and not changes in juice preference, we assessed the waveform stability over time for every neuron. For that, we first extracted the time of the trough and subsequent peak on the average waveform (or alternatively the time of the peak and subsequent trough in inverted waveforms). We then divided all waveforms into 50 bins of equal number and computed the ratio of the amplitude at the average trough time to the amplitude at the average peak time (or vice-versa). This ratio was then z-scored using the mean and standard deviation of the 10 first bins. Neurons whose waveform ratios exceeded ±2.5 sd for 3 consecutive bins were considered unstable. It is important to note that a change in waveform is not an issue per se, so long as the waveform amplitude always stayed above the noise level, which we ensured during our manual spike sorting curation. As expected, rejecting neurons showing changes in waveform amplitudes did not alter the proportion of neurons significantly correlating with local preference (**Fig. 6**, before vs after rejection = 554/1,559 (35.5%) vs 354/1,071 (33%); Chi2<1.12, p>0.29 across areas when comparing before/after rejection). Similarly, the proportion of neurons found to significantly modulate their activity with preference periods (**Fig. 7**) was consistent (before vs after rejection across areas = 430/3,391 (12.7%) vs 246/2,355 (10.45%); Chi2<3.79, p>0.052 across areas when comparing before/after rejection).

#### Statistical procedures

The significance of our results was assessed using permutation tests and multiple correction (using FDR) when appropriate. To assess differences between areas across our measures (e.g., decoding performances or proportion of significant neurons), we fitted generalized linear (or logistic) mixed-effect models (*fitglme* function in Matlab) with area (categorical) as fixed factor as well as monkey (2 levels, categorical) and session (categorical) as random factors (intercept only). When assessing the influence of between-session preference on decoding performance, we used generalized linear mixed-effect models which included an interaction between area and preference (or absolute preference, linear factor) as fixed factor and monkey as random intercept.

### Data and code availability

Behavioral and neurophysiological data will be deposited online upon publication. Code to reproduce the analyses described here will also be made available on our lab GitHub: http://github.com/RudebeckLab/.

## SUPPLEMENTARY MATERIALS

**Supplementary Table 1.**
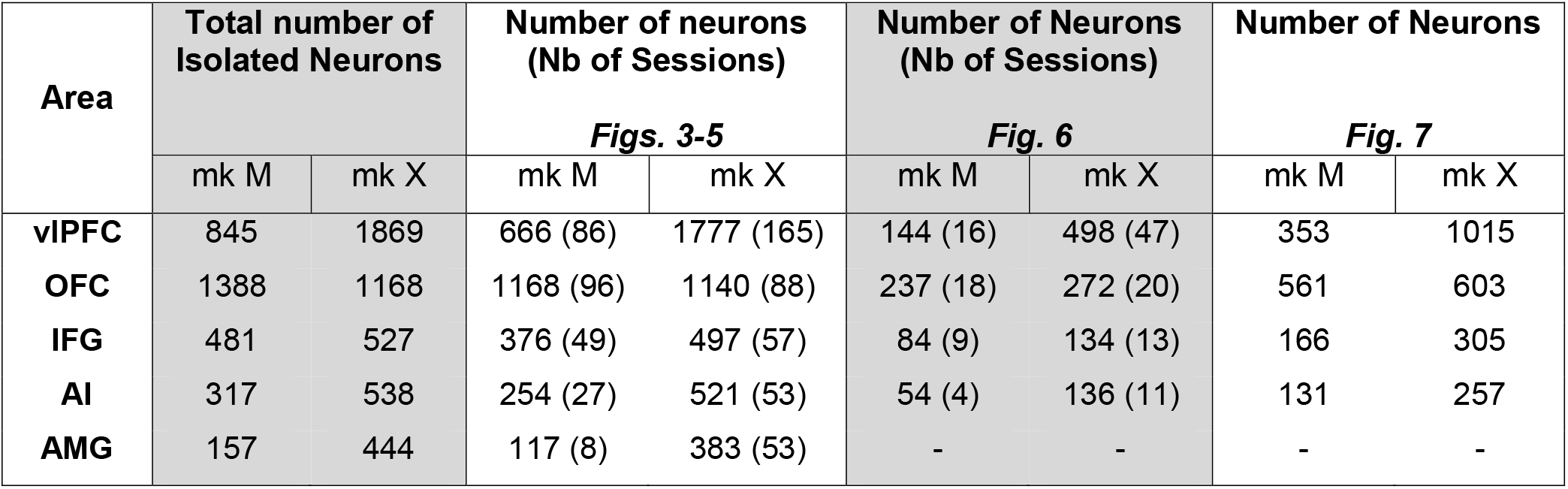
Number of recorded and analyzed neurons for each monkey and main analyses. Numbers varied across analyses if models did not converge or if specific restrictions were used (e.g. Fig. 6 and 7 used a subset of sessions showing significant trend in residuals and preference modulations, respectively). A session was included if 5 or more neurons from the same area were simultaneously recorded.

**Supplementary Table 2.**
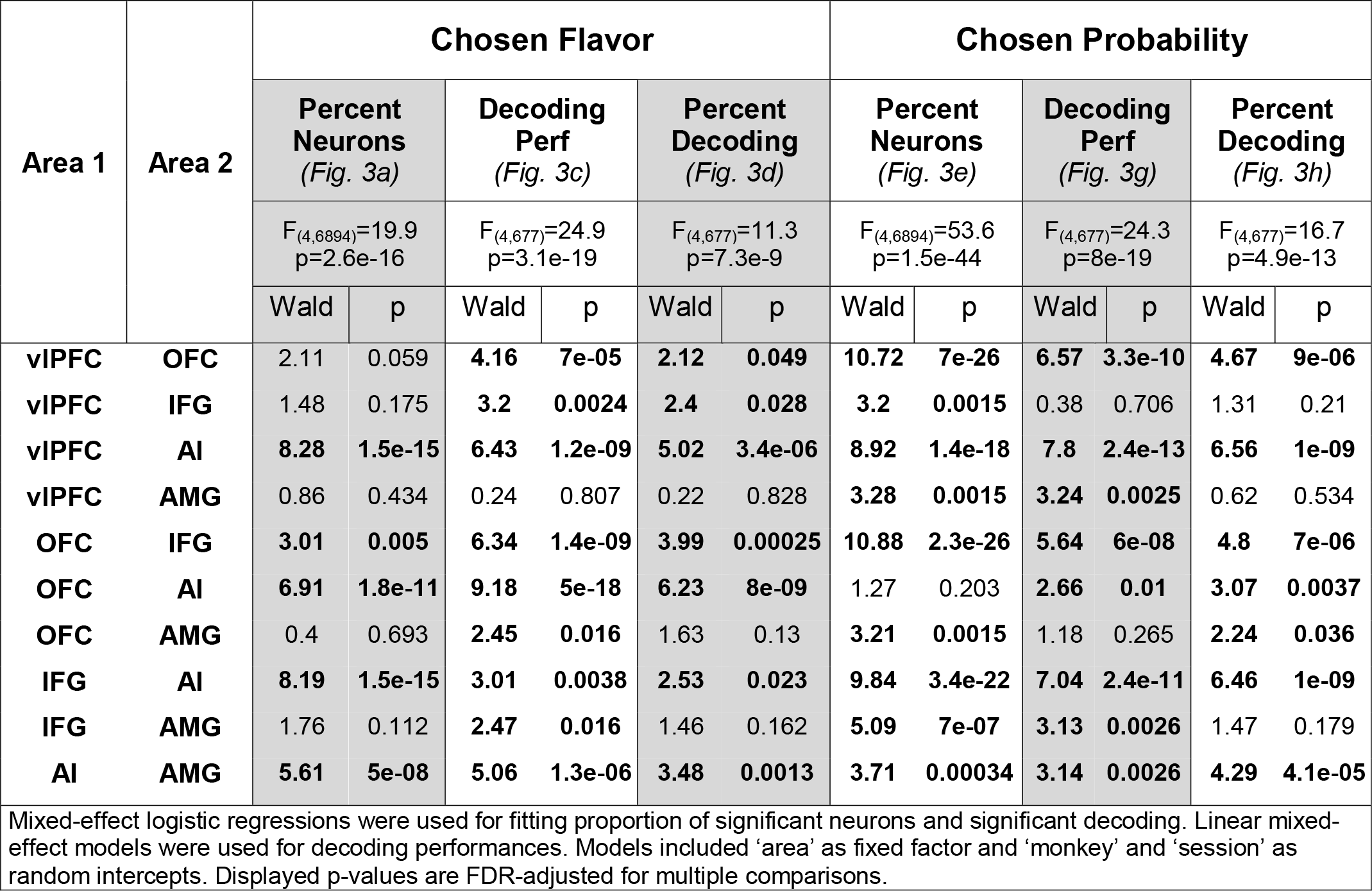
FDR-corrected multiple comparison for area differences in percent of significant neurons and decoding performances for Chosen Flavor and Probability.

**Supplementary Table 3.** FDR-corrected multiple comparison for area differences in percent of significant neurons and decoding performances for Unchosen Probability and Chosen Side.

| Area 1 | Area 2 | Unchosen Probability |  |  |  |  |  | Chosen Side |  |  |  |  |  |
| --- | --- | --- | --- | --- | --- | --- | --- | --- | --- | --- | --- | --- | --- |
|  |  | Percent Neurons<br>(Supp. Fig. 2a) |  | Decoding Perf<br>(Supp. Fig. 2c) |  | Percent Decoding<br>(Supp. Fig. 2d) |  | Percent Neurons<br>(Supp. Fig. 2e) |  | Decoding Perf<br>(Supp. Fig. 2g) |  | Percent Decoding<br>(Supp. Fig. 2h) |  |
|  |  | F<sub>(4,6894)</sub>=68.34<br>p=8.1e-57 |  | F<sub>(4,677)</sub>=15.37<br>p=4.9e-12 |  | F<sub>(4,677)</sub>=8.11<br>p=2.2e-6 |  | F<sub>(4,6894)</sub>=101.7<br>p=2.6e-84 |  | F<sub>(4,677)</sub>=95.4<br>p=2.3e-64 |  | F<sub>(4,677)</sub>=17.3<br>p=1.7e-13 |  |
|  |  | Wald | p | Wald | p | Wald | p | Wald | p | Wald | p | Wald | p |
| viPFC | OFC | 10.19 | 8e-24 | 3.31 | 0.002 | 3.76 | 0.0009 | 9.39 | 1.6e-20 | 7.64 | 1.2e-13 | 4.93 | 3.5e-06 |
| viPFC | IFG | 4.37 | 1.6e-05 | 1.13 | 0.26 | 2.1 | 0.072 | 11.46 | 1.3e-29 | 11.46 | 1.7e-27 | 3.31 | 0.0014 |
| viPFC | AI | 10.78 | 2.2e-26 | 7.09 | 3.5e-11 | 4.65 | 4.1e-05 | 9.73 | 8e-22 | 8.25 | 1.6e-15 | 6.47 | 1.9e-09 |
| viPFC | AMG | 5.18 | 3.9e-07 | 4.4 | 3.9e-05 | 3.01 | 0.009 | 1.33 | 0.185 | 4.08 | 7e-05 | 3.5 | 0.0008 |
| OFC | IFG | 12.03 | 2.7e-32 | 1.58 | 0.127 | 1.07 | 0.355 | 17.74 | 7e-68 | 17.05 | 1.6e-53 | 4.83 | 4.1e-06 |
| OFC | AI | 4.46 | 1.2e-05 | 4.35 | 3.9e-05 | 2.03 | 0.072 | 3.53 | 0.00047 | 2.3 | 0.024 | 2.63 | 0.011 |
| OFC | AMG | 0.86 | 0.388 | 2.05 | 0.059 | 0.01 | 0.995 | 4.31 | 2e-05 | 1.08 | 0.28 | 0.04 | 0.97 |
| IFG | AI | 12.45 | 3.4e-34 | 5.24 | 1.1e-06 | 2.71 | 0.017 | 16.7 | 1e-60 | 16.11 | 6e-49 | 5.66 | 1.1e-07 |
| IFG | AMG | 7.41 | 2.8e-13 | 3.08 | 0.0036 | 0.92 | 0.396 | 9.3 | 2.9e-20 | 11.78 | 1e-28 | 4.62 | 9e-06 |
| AI | AMG | 4.21 | 2.9e-05 | 1.62 | 0.127 | 1.85 | 0.091 | 6.26 | 6e-10 | 2.77 | 0.007 | 2.09 | 0.041 |

Mixed-effect logistic regressions were used for fitting proportion of significant neurons and significant decoding. Linear mixed-effect models were used for decoding performances. Models included 'area' as fixed factor and 'monkey' and 'session' as random intercepts. Displayed p-values are FDR-adjusted for multiple comparisons.

**Supplementary Fig. 1.**
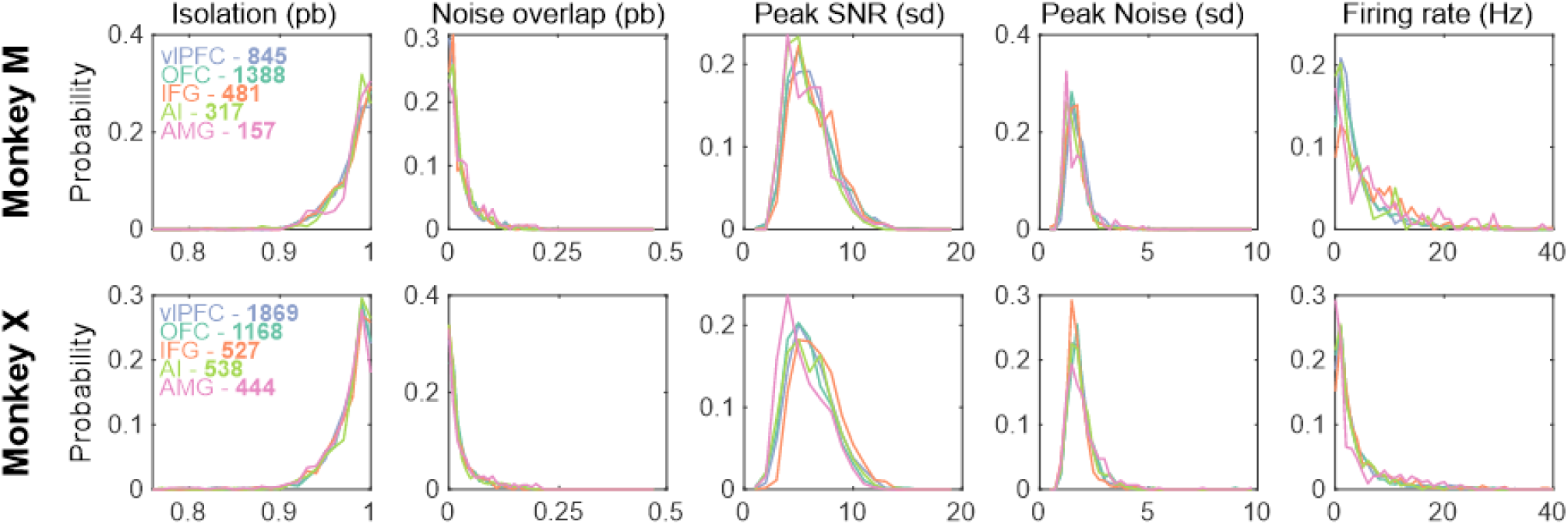
Spike sorting statistics. Distributions for the various estimated metrics extracted during spike sorting with MountainSort for each area in both monkey M (top) and X (bottom). From left to right: isolation quality (probability from 0 to 1; 1 being the purest cluster), noise overlap (probability from 0 to 1; 0 being the least likely to include noise), peak signal to noise ratio (SNR, in sd), peak noise (in sd) and firing rate (in Hz) across the whole session (scaled clipped at 40Hz for display purpose). Note that in cases where two or more clusters were thought to belong to the same putative single neuron during manual curation, the measures associated with that neuron were not recomputed but the reflection of only one of these clusters (e.g., if 2 clusters contained waveforms from the same putative neuron, the isolation measure would be away from 1. Grouping the clusters during curation would not recompute that poor isolation measure). As a result, the estimates shown here are underestimated.

**Supplementary Fig. 2.**
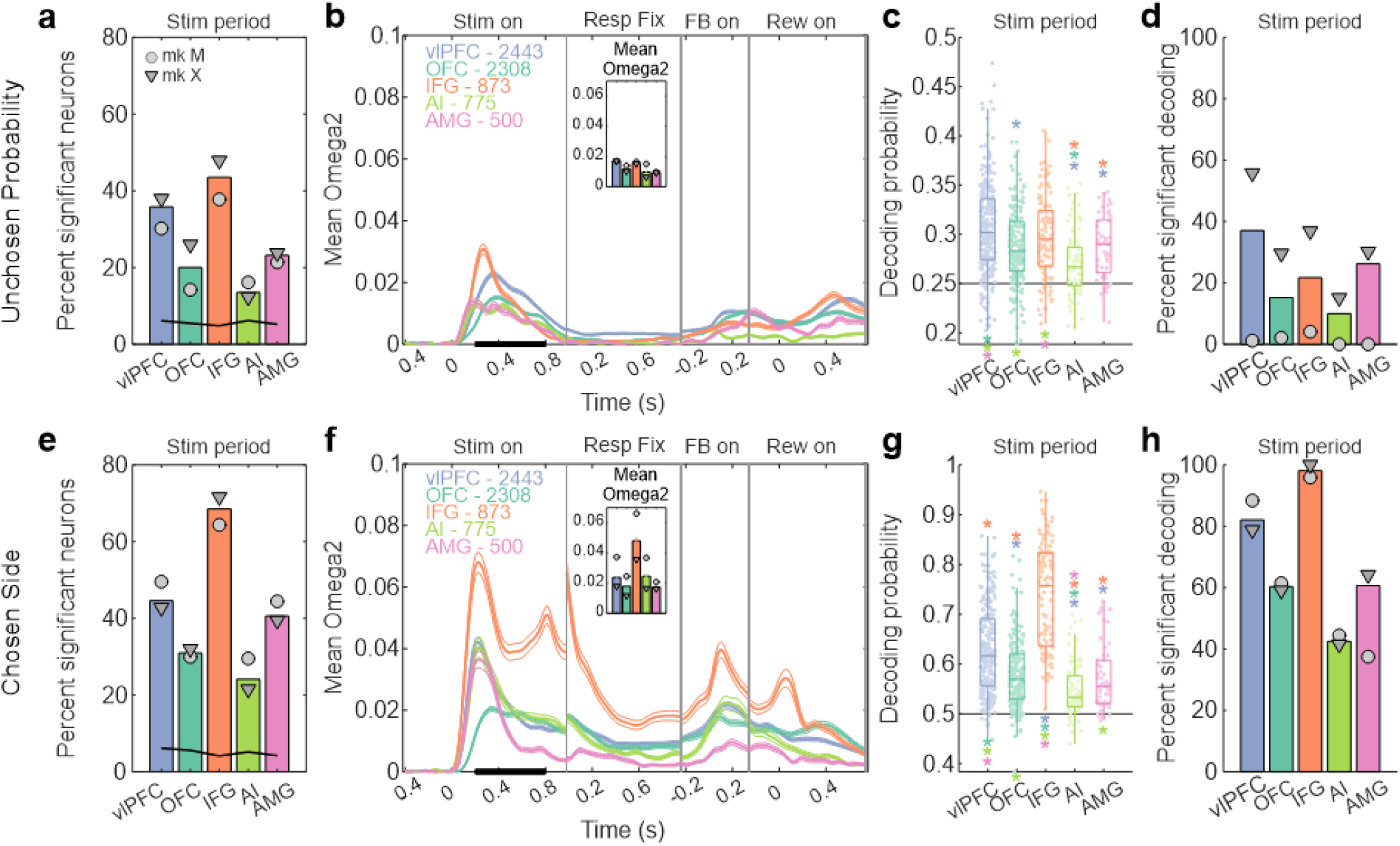
**Unchosen probability (top) and chosen side (bottom) representations** (related to **Fig. 3**). Same as **Fig. 3** but for Unchosen Probability (**a-d**) and Chosen Side (**e-h**) encoding/decoding. (**a, e**) Percent of neurons across areas showing significant firing rate modulation with the parameter of interest during the stimulus period. (**b, f**) Time-resolved average effect size for neurons with significant encoding of the parameter of interest during stimulus period, with inset showing the average effect size during that period. (**c, g**) Decoding performance using simultaneously recorded neurons in each area. (**d, h**) Percent of sessions with a significant decoding performance, assessed using permutation testing.

**Supplementary Fig. 3.**
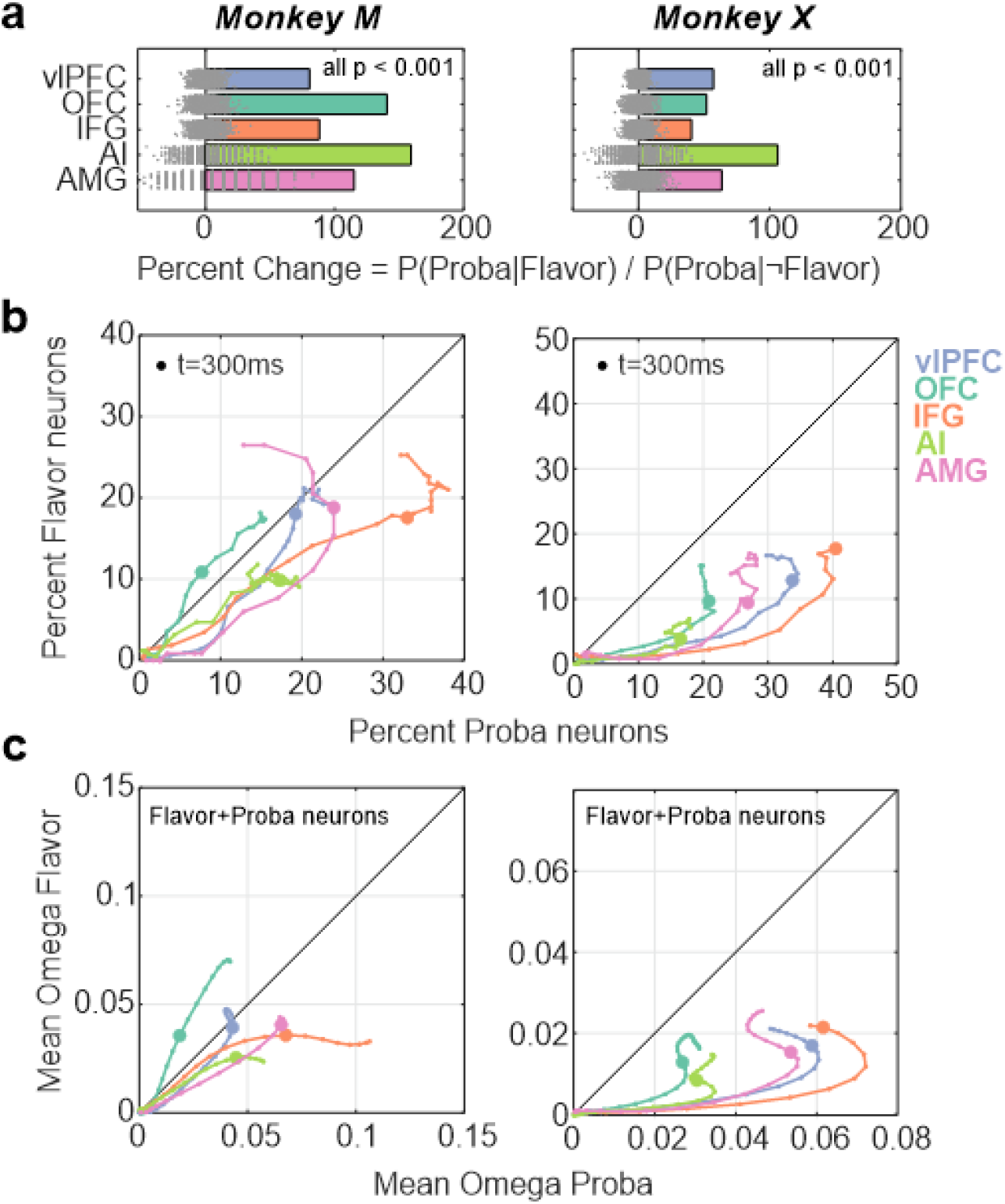
**Mixed selectivity in Flavor and Probability representation in individual monkeys** (related to **Fig. 4**). (**a-c**) Results are shown for monkey M (left) and X (right). (**a**) Estimate of the proportion of chosen flavor neurons also representing chosen probability. Positive values indicate that the same neurons are more likely to encode both parameters, while negative values indicate that different neurons encode the two parameters. Grey dots represent the ratio extracted for each of the 1000 permutations, used to derive significance. (**b**) Evolution over time of the percent of neuron significantly encoding chosen flavor (y-axis) compared to chosen probability (x-axis) for each area. (**c**) Evolution over time of the mean effect size of the significant neurons for flavor and probability.

**Supplementary Fig. 4.**
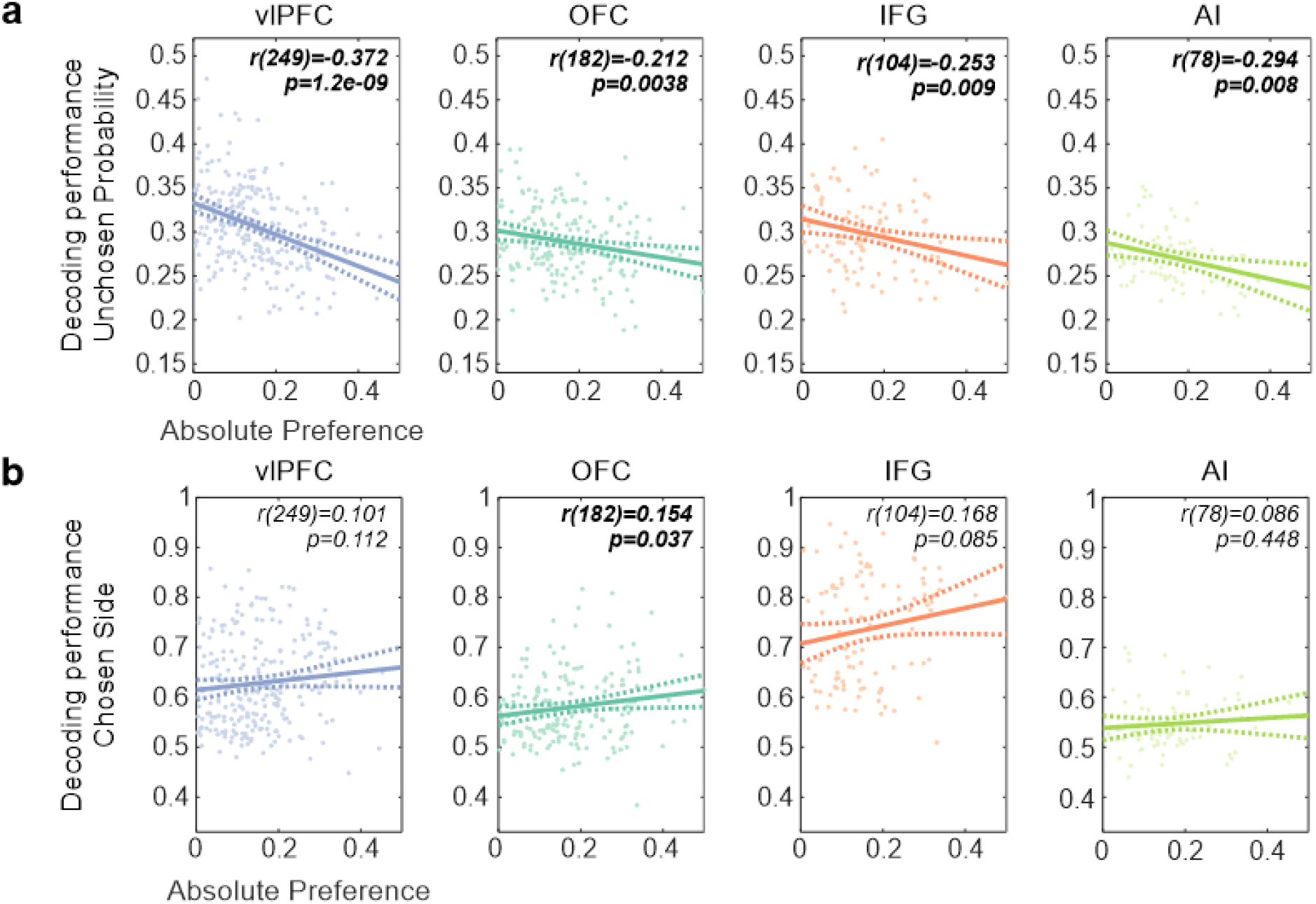
**Unchosen probability and chosen side representation against outcome preference** (related to **Fig. 5**). (**a-b**) Same as **Fig. 5c** but for the unchosen probability (**a**) and the chosen side (**b**). The relation between unchosen probability decoding and preference showed a modest difference between areas (mixed-effect linear regression, interaction abs(preference) x area, F(3,613)=2.7, p=0.046). The representation of the chosen response side did not correlate with preference (mixed-effect linear regression, factor abs(preference), F(3,613)=2.06, p=0.15).

**Supplementary Fig. 5.**
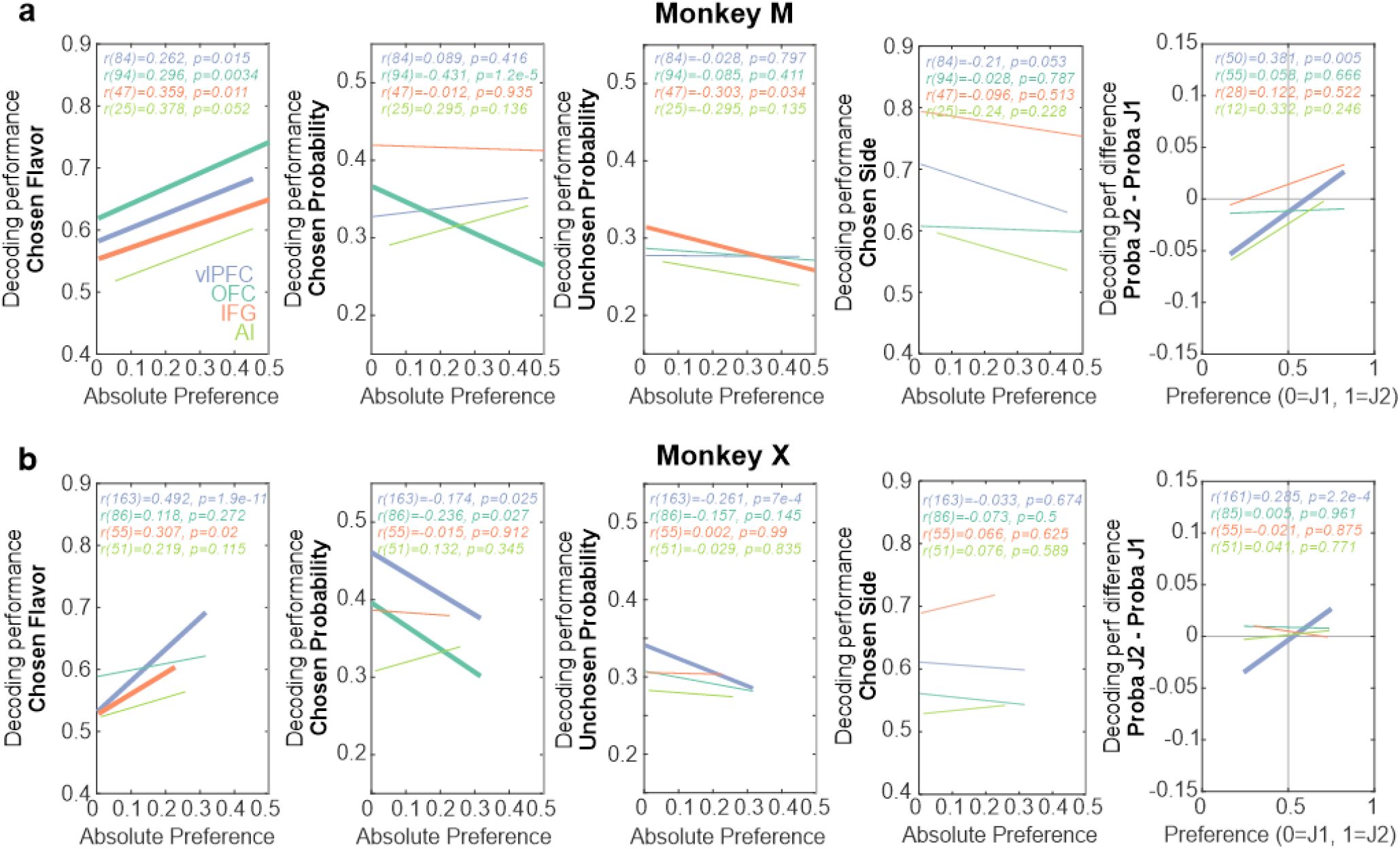
**Outcome flavor and probability representations relate to outcome preference in monkey M (a) and monkey X (b)** (related to **Fig. 5**). (**a-b**) Correlation fits across neuronal populations recorded in monkey M between the decoding performance for outcome flavor, chosen probability, unchosen probability, chosen side and the difference in chosen probability decoding performance when J1 or J2 was chosen (left to right, respectively) and the outcome preference. Bold lines indicate a significant correlation (p<0.05).

**Supplementary Fig. 6.**
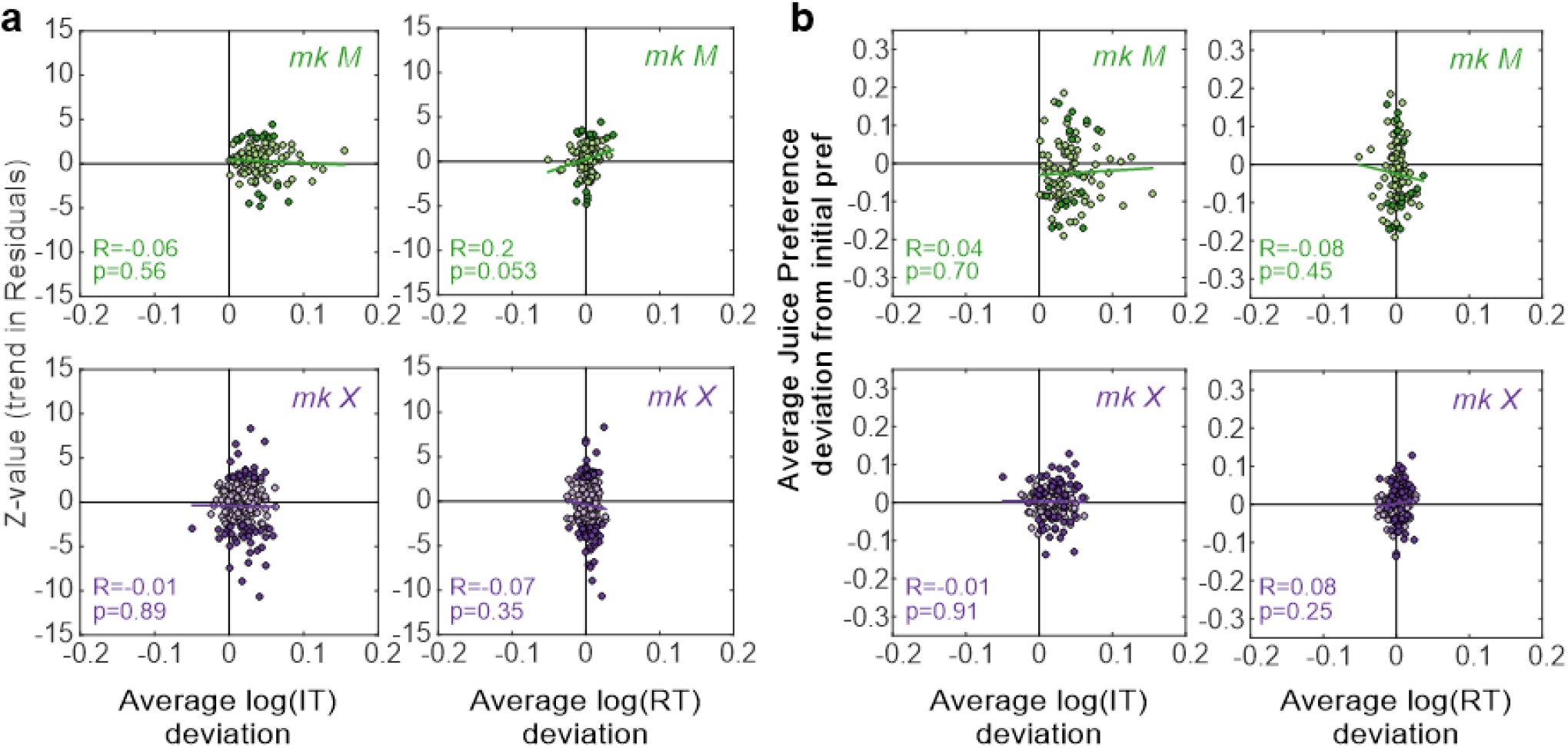
**Absence of between-sessions correlation between response times and Residuals/Preference** (related to **Fig. 6b**). (**a**) Between-sessions trend statistics (Z-value) of the behavioral models’ residuals against the average log(IT) (left panels) and log(RT) (right panels) deviation from the reference period, for monkey M (green, top panels) and X (violet, bottom panels). Each dot represents one session, with the sessions showing a significant trend in residuals highlighted in dark colors (at FDR-corrected p<0.05). (**b**) Same as panel a but showing the average juice preference deviation against log(IT) and log(RT). No significant correlations were observed, highlighting that residuals and juice preference were not associated with motivation changes.

**Supplementary Fig. 7.**
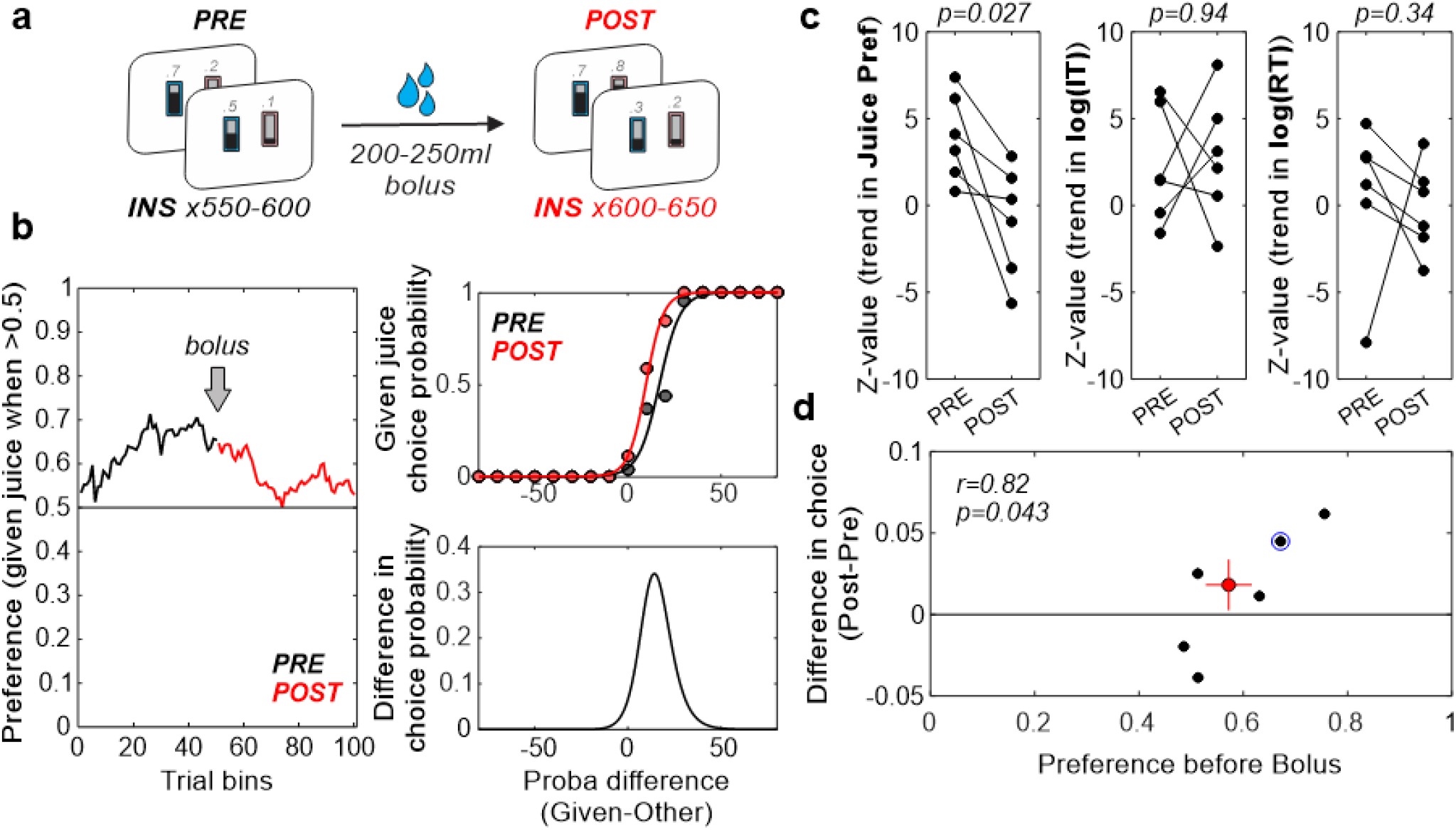
Selective-satiety test in monkey X. (**a**) Schematic representation of the task, where the animal was asked to perform two bouts of Instrumental task trials (PRE and POST periods) separated by the consumption of a large bolus of one of the two available juice. (**b**) Example session showing the preference over trial bins during the PRE (black) and POST (red) periods (left panel), the choice probability for the Given juice against the probability difference for each period (top right panel) and the POST-PRE difference in these choice probabilities (bottom right panel). In this session, the animal slowly developed a preference for one juice, which disappeared once provided with a bolus of that juice. This was also evident on the choice probability, where the animal required greater probability differences to choose the sated juice. (**c**) Z-value statistics across sessions and PRE/POST periods for the trend in juice preference, log(IT) and log(RT) (from left to right panels, respectively) over trial bins. Bolus consumption elicited a consistent decrease in juice preference trends (Wilcoxon signed-rank test; z=2.2, p=0.027), but did not affect log(IT) (z=-0.1, p=0.92) or log(RT) (z=0.94, p=0.34). (**d**) Average difference in choice probability against the average preference before the bolus for each session (black dots). Red dot/lines represent the average/sem across sessions. Blue circle highlights the session shown in panel b. The animal exhibited stronger changes in choice following bolus consumption when their initial preference was stronger (r=0.82, p=0.043).

**Supplementary Fig. 8.**
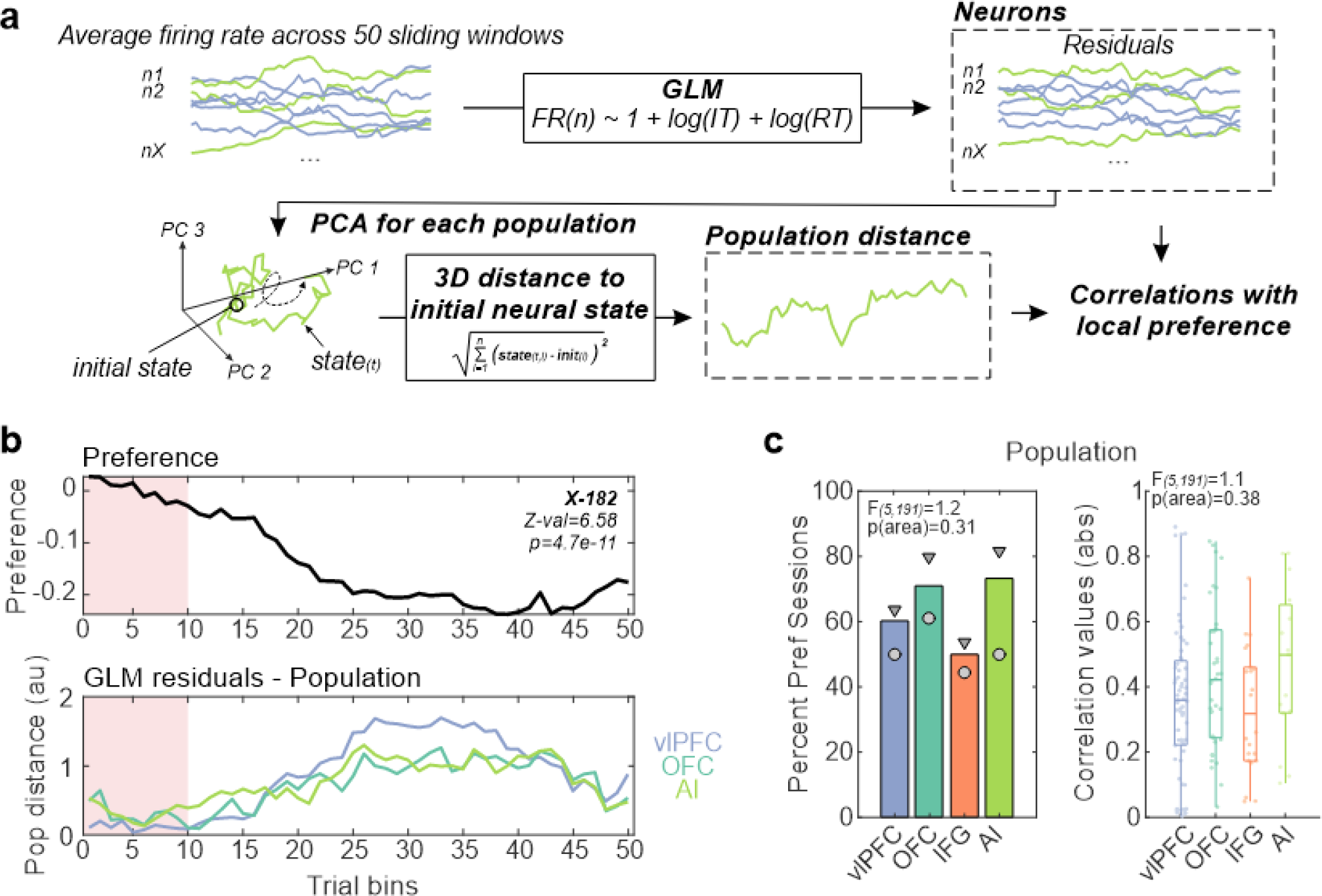
**Analysis pipeline and population modulation with local change in preference** (related to **Fig. 6**). (**a**) Analysis pipeline. In brief, the average firing rate for each neuron was extracted over the 50 sliding windows, before regressing out the influence of log(IT) and log(RT). Models’ residuals were then used to correlate with local preference (neuron level, **Fig. 6e**). Residual activity across population of neurons recorded simultaneously were also reduced to 3 dimensions using PCA, before extracting the Euclidean distance at every trial window from the initial neural state (the average across the first 10 windows). This population distance was then correlated with the local preference (population level, panel c). (**b**) Same example session as **Fig. 6d** but showing the population distance on the bottom panel. (**c**) Same as **Fig. 6e** but for the percent of session and absolute correlation values for population activity distances. Color/marker convention as previously defined. Similar to our neuron level observations, local changes in preference were related to ubiquitous changes in population activity across areas.

**Supplementary Fig. 9.**
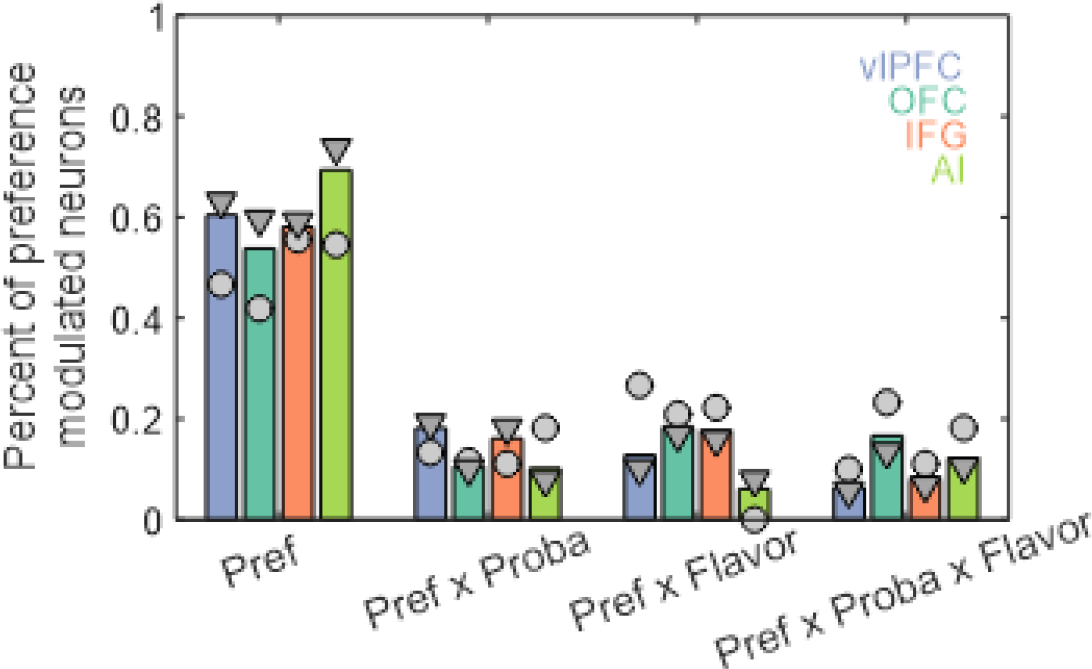
**Relation between preference modulation and outcome flavor/probability** (related to **Fig. 7**). Percent of preference-modulated neurons showing only a main effect of preference (and no interactions), an interaction between preference and outcome probability (with or without main effect of preference), an interaction with outcome flavor or the three-way interaction (with or without “lower” effect) (left to right). Only the OFC population showed as many neurons with a significant main effect as interactions (Chi-2=1.5, p=0.22), whereas other areas showed a greater proportion of main effect only.

